# Hydrogen sulfide modulates plant hypoxic responses through the persulfidation of Plant Cysteine Oxidases

**DOI:** 10.1101/2025.11.05.686772

**Authors:** Y. Telara, S. Akter, A. Aroca, L. Piccigallo, D. Zhang, G. Novi, M. Lavilla-Puerta, D.M. Gunawardana, N. La Monaca, S. Lichtenauer, C. Gotor, M. Schwarzländer, P. Perata, E. Flashman, B. Giuntoli

## Abstract

Hydrogen sulfide (H₂S) is a gaseous molecule historically regarded as toxic. Nevertheless, increasing evidence has brought to light important physiological roles in both animals and plants. In plants, H₂S is involved in environmental and developmental responses, such as stomatal closure and seed germination, and in tolerance mechanisms to different stress conditions like salinity, drought and waterlogging. In this study, we report a function of H₂S as a modulator of hypoxic responses in *Arabidopsis thaliana*. A combination of biochemical and genetic evidence demonstrates that H₂S inhibits the activity of Plant Cysteine Oxidases, the molecular sensors of oxygen, through protein persulfidation to modulate hypoxia-associated responses. Furthermore, we show that H₂S physiology contributes to responses to low oxygen, as disturbing H_2_S production impaired activation of hypoxia-responsive genes and submergence tolerance. Overall, this work introduces H₂S as signalling modulator in plant hypoxic responses and adds a regulatory layer to the plant oxygen-sensing mechanism.

## Introduction

Plants have evolved intricate mechanisms to perceive and respond to low oxygen conditions (hypoxia), which are common in natural environments such as waterlogged or compacted soils (Habibi et al., 2023). A central component of the hypoxic response in *Arabidopsis thaliana* is the Arg N-degron pathway, which coordinates the activity of Ethylene Responsive Factor VII (ERFVII) transcription factors according to O_2_ availability (Gibbs et al., 2011; Licausi et al., 2011). In presence of sufficient O_2_, Plant Cysteine Oxidases (PCOs) catalyse the oxidation of their ERFVII substrates at the level of the N-terminal Cys residue, allowing subsequent ERFVII proteasomal degradation *via* the Arg N-degron pathway (Licausi et al., 2020). Low O_2_ conditions, instead, impair PCO activity. This makes the ERFVII proteins stable, allowing them to migrate into the nucleus and induce the expression of hypoxia-responsive genes (HRGs) (Gibbs et al., 2011; Licausi et al., 2011). The Arabidopsis ERFVII family include five members: Hypoxia-Responsive Element (HRE1 and HRE2) and Related to Apetala2 proteins (RAP2.2, RAP2.3, and RAP2.12) (Licausi et al., 2013). These proteins share a distinctive [MCGGAII(A/S)D] motif, which has been identified as an N-degron (Licausi et al., 2011; Nakano et al., 2006). RAP-type ERFVIIs play a key role in O_2_ sensing, whereas HRE-type ERFVIIs, encoded by hypoxia-inducible genes, rather function downstream in the signalling pathway (Licausi et al., 2010). The ERFVIIs coordinate HRG expression by binding Hypoxia-Responsive Promoter Elements (HRPEs) (Gasch et al., 2016). A core set of HRGs has been identified, across different plant species, that encode proteins involved in hypoxic metabolism, redox regulation, and other processes that remain to be fully characterized (Mustroph et al., 2010; Renziehausen et al., 2024). Hypoxia signalling, however, is not limited to the perception of variations in subcellular O_2_ availability. During O_2_ deprivation, indeed, various signalling molecules accumulate locally, playing a crucial role in coordinating responses to hypoxia. Among them, ethylene, nitric oxide (NO), and reactive oxygen species (ROS) have been extensively studied (Hartman et al., 2019; Sasidharan et al., 2018).

More recently, the gasotransmitter molecule, hydrogen sulfide (H₂S), has been gaining increasing attention within the scientific community. Historically regarded a toxic gas that inhibits respiration, H₂S has emerged in recent years as a signalling molecule in plants and animals (Aroca et al., 2015). In plants, H₂S has been recognized as a key regulator of environmental sensing and developmental programs, including ABA-dependent stomatal closure (Scuffi et al., 2014; Zhang et al., 2020) and seed germination (Sharma et al., 2022). Partially related, a protective role of H_2_S in the biotic and abiotic stress defence response was also suggested (Bhadwal et al., 2024; Xiang et al., 2023; Pantaleno et al., 2024). Notably, pre-treatment with NaHS, a H₂S source, was demonstrated to enhance tolerance to flooding stress and induce the expression of hypoxia-responsive genes in Arabidopsis (Yang et al., 2021). Evidence indicating H₂S accumulation in plant tissues under submergence (Xiao et al., 2020; Yang et al., 2021) highlights its potential involvement in modulating hypoxia-related signalling pathways, a role that remains to be elucidated.

The biosynthesis of H₂S in plants occurs through several pathways; the well-characterized routes are the sulfate (SO₄²⁻) reduction pathway in sulfur assimilation and cysteine metabolism (Saito, 2004). In the former, sulfate is transported into the plastid where its conversion to sulfite (SO₃²⁻) occurs, followed by reduction to sulfide (S²⁻)(Takahashi et al., 2011). Most of the sulfide from the plastid is released as H_2_S and fixed into cysteine, the product of sulfur assimilation, by the enzyme O-acetylserine(thiol)lyase (OASTL), mostly in the cytosol, although other OASTLs are located in the plastid and mitochondria (Hell & Wirtz, 2008). Cysteine participates in the endogenous synthesis of H₂S through the degradation of Cys by various types of cysteine-degrading enzymes (Gotor et al., 2019). Particularly, through the action of L-Cys Desulfhydrases (LCDESs), it leads to the release of H₂S, pyruvate and ammonia, and in Arabidopsis, one cytosolic LCDES, named DES1, has been characterized in detail (Álvarez et al., 2010). Cys catabolism plays a crucial role when plants are exposed to oxidative stress or O_2_ deprivation. Under these stresses, Cys catabolism is upregulated, leading to increased H₂S production, which can act as a signalling molecule to mitigate the effects of the stress (Yang et al., 2021).

While the precise mechanisms of H₂S signalling in plants are yet to be completely understood, one of the proposed modes of action involves protein persulfidation (Aroca et al., 2015; Mustafa et al., 2009). This post-translational modification occurs on the thiol groups (-SH) of Cys residues, converting them into persulfides (-SSH) (Aroca et al., 2017; Aroca et al., 2018). However, the direct interaction between H₂S and Cys thiols is thermodynamically disfavoured (Cuevasanta et al., 2015; Filipovic et al., 2018). Therefore, persulfidation typically requires the prior oxidation of thiols to sulfenic acids (-SOH) or S-nitrosylation to nitrothiols (-SNO) *via* ROS or NO, respectively (Vignane & Filipovic, 2023). This adds a layer of complexity into the biochemistry of persulfidation, suggesting a finely tuned regulatory mechanism involving multiple signalling molecules, with significant consequences for protein function (Aroca et al., 2021; Aroca et al., 2017). Persulfidation is conserved across all domains of life and plays a key role in regulating the localization, conformation and activity of numerous proteins, with profound effects on cellular regulation (Aroca et al., 2018). A proteomic analysis revealed that, out of over 900 proteins in Arabidopsis susceptible to S-nitrosylation, more than 600 also represent persulfidation targets (Aroca et al., 2017). Among these dual targets, Plant Cysteine Oxidase 3 (PCO3) was identified, which is particularly notable in the context of hypoxia. In this study, we investigate the role of H₂S in the response to hypoxia in Arabidopsis. Using biochemical and genetic analyses, we observed that H₂S directly modulates the activity of PCO enzymes, influencing the ability of plants to sense O_2_.

## Results

### Sulfate burst stimulates ERFVII-mediated responses in aerobic conditions

Sulfate metabolism is the route of H₂S biosynthesis in plant cells; therefore, we speculated that modulating this process could influence hypoxia responses of Arabidopsis seedlings. We sought to stimulate a transient burst in H_2_S production through the reductive sulfate assimilation pathway by modifying S availability in the nutrient solution. Col-0 seedlings grown for 5 days under S deficiency (S-, 25 µM MgSO₄) were transferred to optimal S conditions (S+, 750 µM MgSO₄) for 6 hours (**Fig. 1a)** and the effects of S re-supply on marker gene expression was evaluated. Seedlings grown under low S for 5 days showed signs of starvation, as revealed by elevated expression of the S-deficiency markers *SULFATE TRANSPORTERS (SULTR1;1* and *SULTR4;2)* in comparison with plants grown for the same time on S-replete media (**Fig. 1b**). Their expression decreased after S re-integration, suggesting the restoration of a normal S status. On the contrary, hypoxia marker genes were not elevated during S depletion, but three out of the four measured markers were significantly induced after S-reintegration (specifically, *ADH1*, *PCO1*, and *PDC1*) (**Fig. 1b**). The induction of these genes in fully aerated seedlings indicates that low O_2_ signalling can be modulated by S assimilation. A meta-analysis of transcriptomic data from an existing study of S re-integration in *A*. *thaliana* (Bielecka et al., 2015) showed, compatibly, the induction of several HRGs already 30 minutes after sulfate resupply, including *HRA1*, *PDC1*, *PCO1* and *WIP4* measured here (**Suppl. Table 1**).

**Figure 1.**
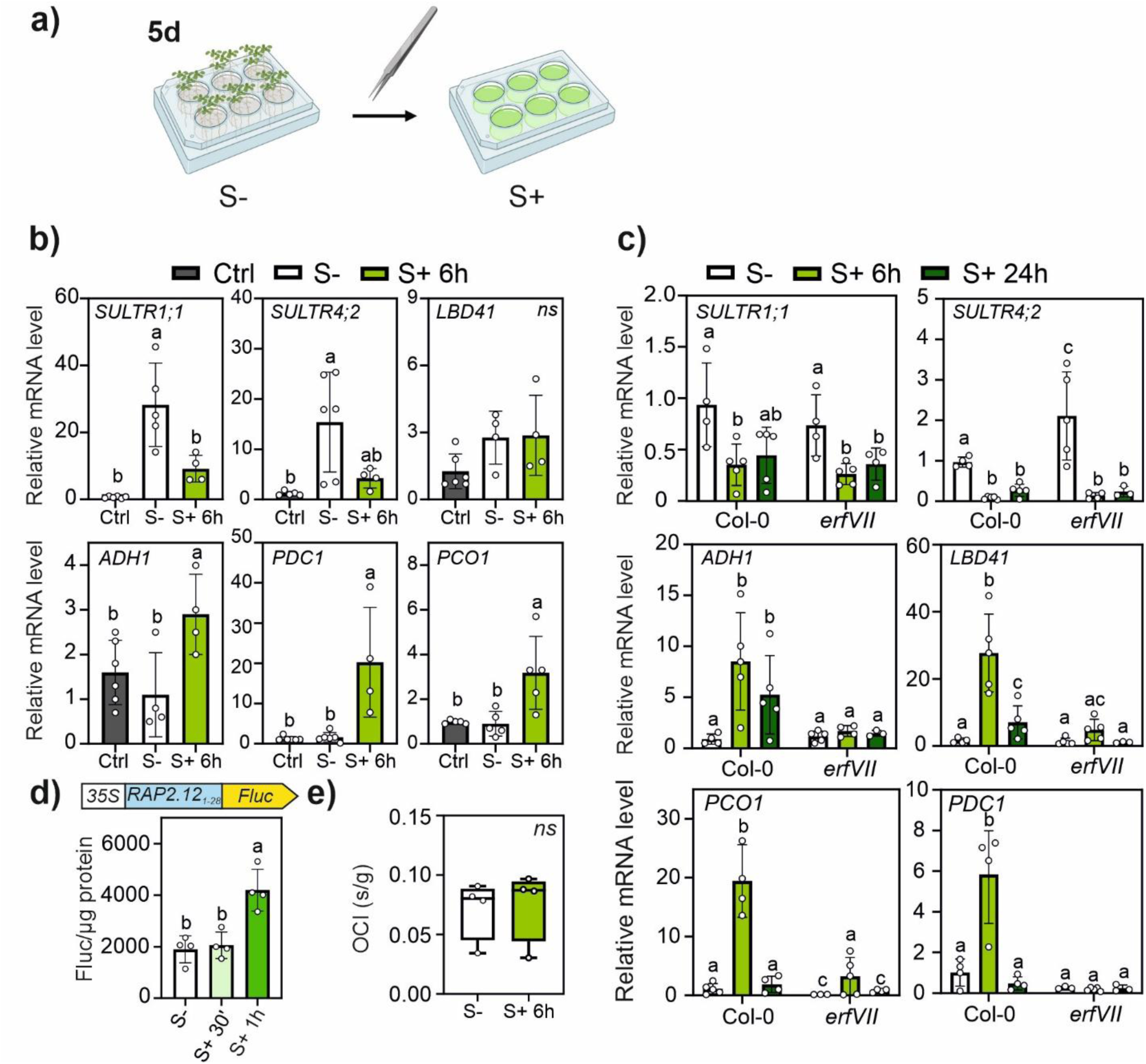
Effects of S-resupply on the responses to hypoxia and O_2_ consumption. **a)** Schematic representation of the S re-integration experiments: seedlings were sampled after 5 days of growth on S deficiency (S-, 25 µM) or 6 h after S re-integration (S+, 750 µM) in the liquid media. S was provided as MgSO_4_. **b)** Gene expression in Col-0 seedlings treated as in (a). Data (mean ± SD, n=5) are relative to a control sample growth for 5 days under S optimal condition (“Ctrl”, 750 µM S). **c)** Gene expression in Col-0 or *erfVII* mutant treated as in (a), after 6-or 24-hours re-integration. Data (mean ± SD, n=5) are relative to one S-sample. Letters indicate statistically significant differences among conditions and genotypes after two-way ANOVA and Tukey-Kramer post-hoc test (p<0.05). **d)** O_2_ consumption by Col-0 seedlings treated as in (a) and transferred to closed vials equipped with a fluorescent O_2_-sensitive spot (see Materials and Methods). OCI, Oxygen Consumption Indicator (see main text). **e)** Reporter activity in *35S:RAP2.12_1-28_:Fluc* (*28RAPFluc*) seedlings treated as in (a). Firefly luciferase activity was measured before the transfer and after 30 minutes and 1 hour from the recovery and normalised to total soluble proteins (Fluc/µg total proteins). Data are mean ± SD (n=4). Letters in (b) and (d) indicate statistically significant differences between different conditions after one-way ANOVA and Tukey-Kramer post-hoc test (p<0.05).

To understand the role of the ERFVIIs in this phenomenon, we compared the response to 6- or 24-hour S re-supply in aerobic seedlings of a pentuple *erfVII* mutant (Abbas, Berckhan, Rooney, Gibbs, Vicente Conde, et al., 2015) with the wild-type. S-deficiency markers had comparable expression profiles in both genotypes, indicating no involvement of the ERFVIIs in their regulation in response to variations in S provision (**Fig. 1c**). S-resupply for 6 hours was again associated with the induction of the hypoxia markers in Col-0. The expression of *PCO1, PDC1* and *LDB41* reverted to basal levels by 24 hours, whereas *ADH1* was still induced. In contrast, the *erfVII* mutant showed no induction of the hypoxia markers, except for a marginal response of *PCO1* (**Fig. 1c**). These results indicate that the transition from S deficiency to optimal S availability, under aerobic conditions, triggers HRG expression in an ERFVII-dependent manner. To further support the role of ERFVII in this response, we used a *35S*:*RAP2.12_1-28_:Fluc* reporter line of Arabidopsis (hereafter, *28RAPFluc*), to monitor the activity of the Cys N-degron pathway (Weits et al., 2014). This translational fusion links the luminescence signal to the N-terminal modifications of RAP2.12, while excluding any regulation impinging on downstream domains of the protein such as phosphorylation (Kunkowska et al., 2023) or SINAT-dependent proteolysis (Papdi et al., 2015). S re-supply stimulated reporter activity (**Fig. 1d**), indicating that, consistent with the observed HRG upregulation, the S burst promoted RAP2.12_1-28_ stabilization under aerobic conditions through modulation of the Cys N-degron pathway. Re-integration of phosphate or iron, after growth on the corresponding deficient media, had no effects on *28RAPFluc* output **(Suppl. Fig. 1)**.

We next considered the possibility that the anaerobic response induced by S deficiency/re-supply was triggered by regulation of metabolism and respiration (Martin & Maricle, 2015; Pietri et al., 2011). Specifically, we tested whether S re-supplementation might induce a rapid and intense increase in O_2_ consumption, thereby generating local hypoxia and inducing anaerobic responses. We thus measured the O_2_ consumption rate in the same conditions as those in **Fig. 1a**. Following the treatments, seedlings were immersed in water in sealed vials, where the decline in dissolved O_2_ due to plant metabolism could be monitored over time thanks to a fluorescent O₂-sensitive sensor (oxyspot). To enable data comparison, we expressed O_2_ consumption as OCI (Oxygen Consumption Indicator), which represents the time required for oxygen saturation to decrease from 85% to 70% of the initial value (full saturation under 21% O_2_ atmosphere), normalized to fresh weight. OCI was unchanged after 6 hours S re-integration, suggesting that flux through respiratory metabolism is not affected to any extent by the treatment (**Fig. 1e**). A control treatment with 8 mM pyruvate, which has previously been observed to induce a higher respiratory rate (Zabalza et al., 2009), resulted instead in increased OCI (**Suppl. Fig. 2a**), however this was not associated with changes in HRG expression (**Suppl. Fig. 2b-c**). Together, the data confirm that the S-induced hypoxic response observed in **Fig. 1a-b** was not caused by altered respiration rates.

### Hypoxic responses generated by sulfate re-integration are linked to H_2_S

Aiming to verify that H₂S was involved in the normoxic induction of the hypoxic response, we used NaHS as a sulfide donor. *RAP28Fluc* seedlings exposed to different concentrations of NaHS for 2 hours showed that H₂S-mediated RAP2.12_1-28_ stabilization dose-dependent and non-linear: 50 µM NaHS treatment had no significant effects, whereas concentrations ranging from 100 µM to 1 mM caused reporter stabilization to a similar extent, as compared to untreated samples (**Fig. 2a**). We then investigated the temporal dynamics of *RAP28Fluc* response to 100 µM NaHS treatment (the lowest dose associated with an effect). Luciferase activity was significantly increased after 1 and 2 hours, returning to control levels after 4 hours (**Fig. 2b**). A similar transient response was observed upon treatment with 500 µM NaHS (**Suppl. Fig. 3a**). Higher luciferase activity in the reporter lines after 2 hours NaHS treatment was compatible with RAP2.3-HA protein stabilization in a *35S:RAP2.3^3xHA^* transgenic line (Gibbs et al., 2014) (**Suppl. Fig. 3b**).

**Figure 2.**
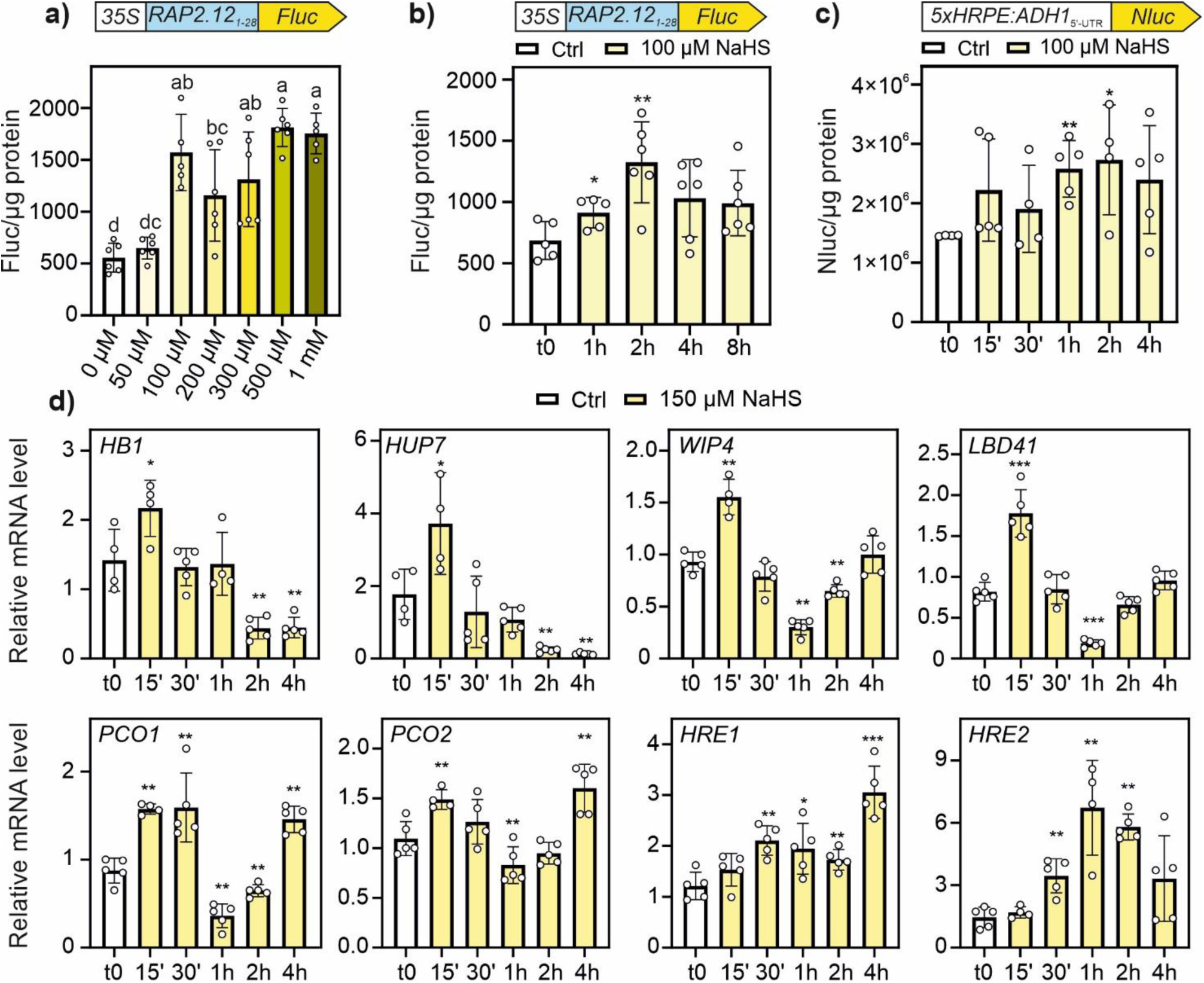
Characterization of the hypoxia-like response to H_2_S supplementation. **a)** Luciferase activity in 7-days old *35S:RAP2.12_1-28_:Fluc* (*28RAPFluc*) seedlings treated with different concentrations of NaHS for 2 hours (n=5). Different letters indicate statistically significant differences according to one-way ANOVA followed by Tukey’s post hoc test (p<0.05). **b)** Luciferase signal in 7-days old *28RAPFluc* seedlings in a time-course experiment with 100 µM NaHS (n=6). **c)** Nanoluciferase activity in of 7-days old *HRPE:Nluc* seedlings in a time-course experiment with 100 µM NaHS (n=5). **d)** Hypoxic marker gene expression in 7-days old Col-0 seedlings at different timepoints after the supplementation of 150 µM NaHS. All data are shown as mean ± SD. Data in (a-c) were normalised with total proteins in the extracts. Asterisks indicate statistically significant differences between treated and t_0_ samples (b-d), or treated and control samples (a) after Student’s t-test (* p<0.05; ** p<0.01; *** p<0.001).

To verify whether, alongside ERFVII stabilization, H₂S also triggered the induction of hypoxia marker transcripts, we made use of the transcriptional reporter line *HRPE:Nluc* of Arabidopsis (Akter et al., 2024), in which *NANOLUCIFERASE* expression is driven by a synthetic *5xHRPE:ADH1*_5’-UTR_ promoter (Akter et al., 2024). Following treatment with 100 µM NaHS, the Nluc signal increased significantly after 1 hour, remained stable up to 2 hours (**Fig. 2c**). Treating *HRPE:Nluc* seedlings with 500 µM NaHS resulted in a more pronounced and faster response (**Suppl. Fig. 3c**).

Signal reversal in the previous assays might have been partially concealed by the stability of the luciferase proteins (Urquiza-García et al., 2019) leading us to turn to mRNA assessment to complement the above observations. Profiling of endogenous gene expression changes at different NaHS treatment levels showed the transient and dose-responsive signature of the hypoxia-like response. A set of hypoxic genes composed by *HB1*, *HUP7*, *WIP4*, and *LBD41* was rapidly induced by NaHS supplementation (as early as 15 minutes after treatment), reversed to baseline levels within 30 minutes and underwent inhibition afterwards (**Fig. 2d**). This behaviour was consistently observed in a treatment range from 150 to 500 µM, whereas any transcriptional response was barely visible at 100 µM (**Suppl. Fig. 4**). *PCO1* and *PCO2*, showed a biphasic response characterized by later re-induction, after rapid and more stable induction initial induction followed by repression (**Fig. 2d**). A slower response was, instead, observed for *HRE1* and *HRE2* (**Fig. 2d**). Again, the same profiles were observed across a range of NaHS treatments (**Suppl. Fig. 5**). Finally, *HRA1*, *PDC1* and *ADH1* were unaffected by NaHS, except for slight induction of *ADH1* at 500 µM (**Suppl. Fig. 6**). Altogether, these results suggest that H₂S is associated with a rapid and reversible modulation of the activity of proteins involved in the Cys N-degron pathway, causing distinct changes in downstream transcription.

### PCO enzymatic activity is impaired by H_2_S

ERFVII stabilization and HRG triggering relies on the inactivation of the PCO-initiated Cys N-degron pathway (Gibbs et al., 2011; Lavilla-Puerta et al., 2023; Licausi et al., 2011). Given the swift dynamics of the hypoxic response observed during NaHS treatment, we hypothesized that H₂S may directly affect PCO activity. To test this, we performed an *in vitro* assay to examine the role of H₂S in modulating the activity of recombinant *At*PCO4, as a representative member of the PCO family (White et al., 2018). The activity of *At*PCO4 was monitored by examining Nt-Cys oxidation in a RAP2_2-15_ peptide in the presence of O_2_ (White et al., 2017). *At*PCO4 was at first pre-incubated for 10 minutes with or without 1 mM NaHS or with 1 mM H₂O₂, a known inhibitor of PCO activity (Akter et al., 2024). The data showed that H_2_S inhibits *At*PCO4 activity (**Fig. 3a**). Pre-incubation periods ranging from 5 to 30 minutes were then tested, to assess the effect of NaHS exposure duration on *At*PCO4. A progressive reduction in RAP2_2-15_ oxidation was observed with longer pre-incubations, indicating that H_2_S inhibits *At*PCO4 activity in a time-dependent manner (**Fig. 3b**). When the dose-response effect was examined, an “apparent’’ IC_50_ value of 1092 μM was determined, albeit the data does not completely reflect a competitive inhibition model, potentially due to a persulfidation effect (**Fig. 3c**). We next examined whether reducing agents could restore the enzymatic activity of H_2_S-inhibited *At*PCO4 through reversal of persulfidation. Recombinant *At*PCO4 was initially treated with 5 mM NaHS for 30 minutes, followed by incubation either with the biological reducing agent 10 mM glutathione (GSH) or 10 mM tris(2-carboxyethyl)phosphine (TCEP) for varying durations (1, 5, 15, or 30 minutes), before exposure to the RAP2.12 substrate. Both reducing agents partially restored *At*PCO4 activity in a time-dependent manner; after 30 minutes of treatment, TCEP recovered approximately 20% of the enzyme’s maximal activity, whereas GSH restored about 14% (**Fig. 3d-e**). To assess whether H₂S-mediated inhibition is a general feature of the Arabidopsis PCO family, we extended our analysis to all five isoforms under two conditions: (i) pre-incubation with 10 mM GSH followed by 5 mM NaHS, and (ii) NaHS treatment followed by GSH incubation. In both experimental setups, partial restoration of activity was observed for all PCO isoforms, with the most substantial recovery occurring in PCO1 and PCO5 (**Fig. 3f**). Although the reducing potential of GSH can be variable *in vitro* (due to trace amounts of oxidised GSH), these results support a redox-sensitive and partially reversible mechanism of H₂S-mediated inhibition which likely involves one or more of the Cys residues in PCOs.

**Figure 3.**
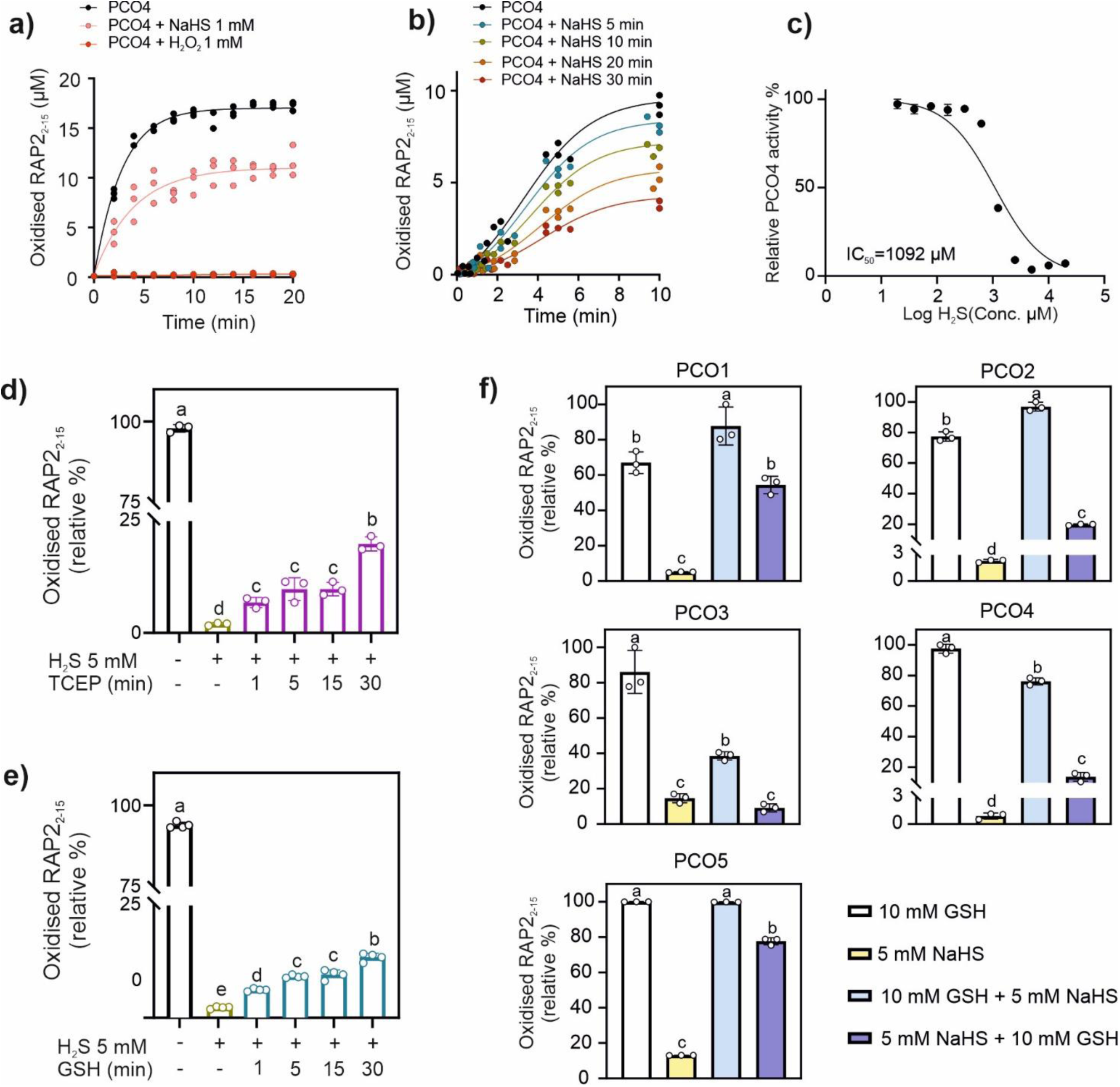
Modulation of PCOs activity *in vitro* induced by H_2_S exposure. **a)** H_2_S inhibitory effects on *At*PCO4 enzymatic activity *in vitro* measured as oxidation of RAP2_2-15_ peptide. The enzyme was pre-incubated for 10 minutes with the indicated amount of NaHS, or with H_2_O_2_, before the assays. Non-treated *At*PCO4 and 1 mM H_2_O_2_ were included to compare the effect of H_2_S on *At*PCO4 activity (n=3). **b)** Effect of different exposure times (5, 10, 20, or 30 min) to 5 mM NaHS on *At*PCO4 activity (n=3) under the same conditions of a). **c)** Dose-response curve for NaHS treatments of 2 µM *At*PCO4 activity; reactions were performed exposing *At*PCO4 for 30 minutes to NaHS ranging from 20 μM to 20 mM. Data are mean ± SD (n = 3). **d)** Recovery of *At*PCO4 enzymatic activity induced by glutathione (GSH) and **e)** tris(2-carboxyethyl)phosphine (TCEP). Data are mean ± SD (n=3-4). In (d) and (e) letters indicate significant differences between treatments after one-way ANOVA followed by Tukey HSD test (p<0.05). **f)** H_2_S-mediated inhibition of *At*PCOs and their recovery of enzymatic activity after 30 minutes pre-incubation with 5 mM NaHS, in presence or absence of 5 mM GSH (n=3). Data are mean ± SD (n = 3). Letters indicate statistical differences evaluated using One-way ANOVA followed by Tukey HSD test (p<0.05).

### AtPCO4 is modulated by H_2_S through persulfidation

H₂S has been proposed to regulate protein function through persulfidation, a redox-sensitive post-translational modification (PTM) in which Cys thiol groups (-SH) are converted to persulfides (-SSH) (Aroca et al., 2015; Mustafa et al., 2009). Given the observed sensitivity of PCO enzymatic activity to H₂S treatment **(Fig. 3d)**, we investigated the potential interaction between H₂S and *At*PCO4 Cys residues. We examined the effect of H₂S on cysteine modification of *At*PCO4 using the BioDiaAlk probe, which selectively reacts with cysteine-derived sulfinic acid moieties (Akter et al., 2018). *At*PCO4 was treated with 10 mM NaHS, either alone or in combination with 1 mM H₂O₂, followed by BioDiaAlk labeling. In the presence of H₂O₂ alone, a marked increase in sulfinylation of *At*PCO4 was observed, indicating the formation of sulfinic acid. However, when H₂S was co-applied with H₂O₂, sulfinylation signals were substantially reduced or completely absent (**Fig. 4a** and **Suppl. Fig. 7**). These results imply that H₂S directly interacts with oxidized Cys residues on *At*PCO4, potentially preventing their further oxidation to sulfinic acid.

**Figure 4.**
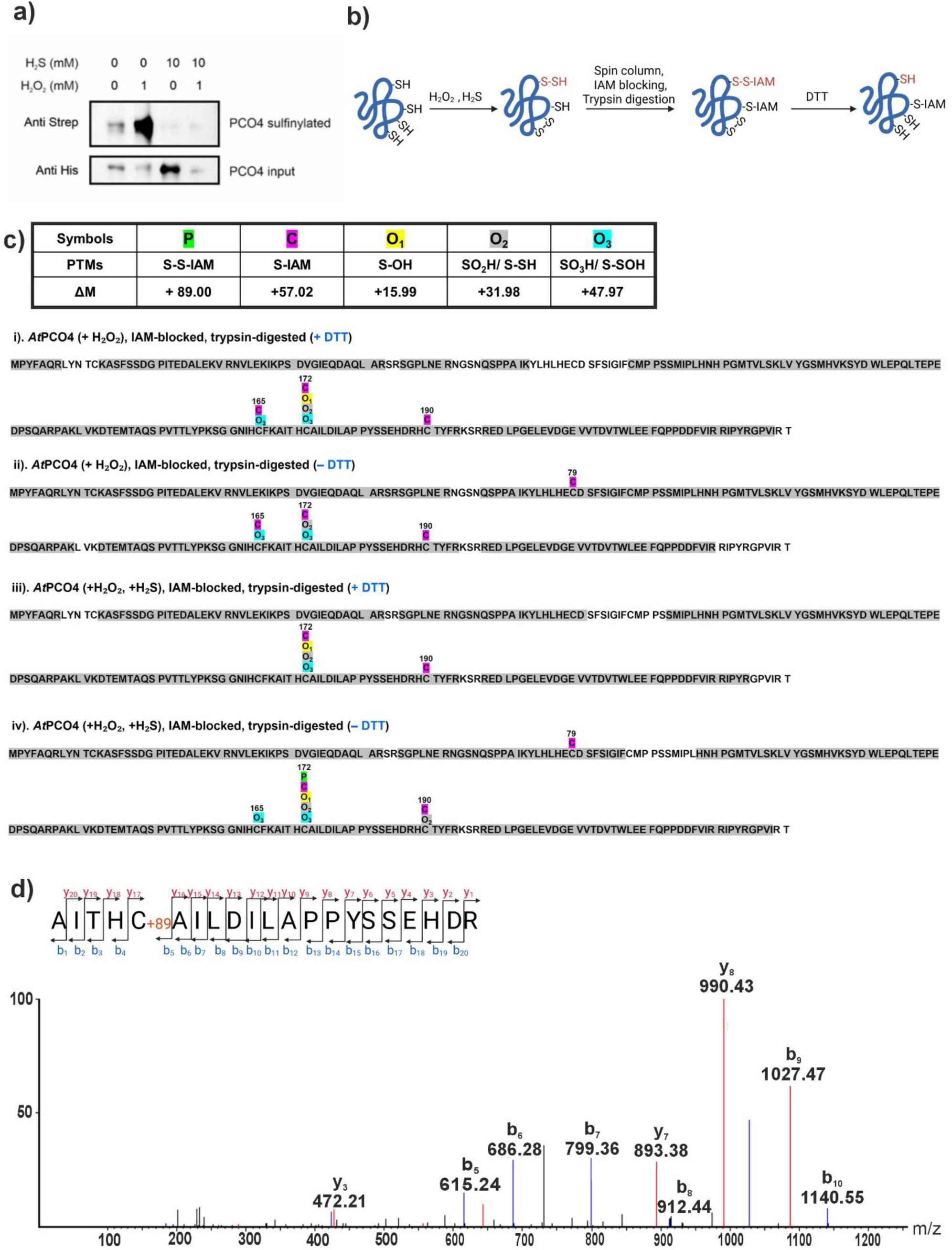
Detection of persulfidation in *At*PCO4 by differential alkylation-mass spectrometry strategy. **a)** Detection of *At*PCO4 sulfinylation *in vitro*. Purified *At*PCO4 was treated in combination with 10 mM H_2_S and/or 1 mM H_2_O_2_ and the amount of *At*PCO4 sulfinylated detected using the BioDiaAlk probe specific for sulfinic acid (Akter et al., 2018). The full-size images of the membranes can be found in Suppl. Fig. 8. **b)** Schematic representation of the differential IAM labeling workflow used for persulfidation detection. Proteins were first treated with NaHS (H₂S) to induce persulfidation, either alone or in combination with H₂O₂. Free thiols and persulfidated cysteines were then alkylated with iodoacetamide (IAM), blocking accessible –SH and –SSH groups. Following tryptic digestion, peptides were incubated with or without dithiothreitol (DTT) to reduce persulfides or other reversible oxidative modifications (e.g., disulfides). By comparing the thiol content in DTT-treated and untreated samples, putatively persulfidated cysteine residues were identified. **c)** Summary of potential cysteine modifications and corresponding mass shifts in H₂O₂ + NaHS-treated *At*PCO4 (top), and *At*PCO4 peptide sequence coverage after MS/MS analysis. Modifications observed at specific cysteine residues following DTT or no-DTT treatment are annotated. **d)** LC-MS/MS spectrum showing the fragmentation pattern of the modified peptide, including annotated b-and y-ions of interest.

As all five *At*PCOs were inhibited by H_2_S treatment, we looked for conserved Cys residues that may act as potential PTM targets. Cys^12^, Cys^79^, Cys^88^, Cys^172^, and Cys^190^ (*At*PCO4-1 numbering) (Dirr et al., 2025) are conserved across all isoforms, and an additional cysteine residue (Cys^165^) is present in all isoforms except *At*PCO1 (**Suppl. Fig. 8a**). To investigate which of those residues were persulfidated by H_2_S treatment, we performed LC-MS/MS–based proteomic analysis on NaHS-treated and untreated *At*PCO4 samples, adapting a differential alkylation-based method for persulfidation detection, following the principles described by Aroca et al. (2015) and Zivanovic et al. (2019). In this approach, both free thiols and persulfidated cysteines were initially alkylated with iodoacetamide (IAM). After tryptic digestion, the peptide mixture was split: one aliquot was subjected to DTT reduction, selectively releasing persulfide-linked IAM adducts, while the other was left untreated **(Fig. 4b)**. While more than one Cys residue was found to be oxidised or modified by IAM, under the conditions tested, Cys^172^ of *At*PCO4 was the only residue identified by MS/MS to undergo persulfidation in response to H₂S **(Fig. 4c)**. In the non-reduced (–DTT) sample, persulfidated cysteines retained an additional +89 Da mass shift (comprising +32 Da from S and +57 Da from IAM). In contrast, the DTT-treated sample showed conversion of these sites into reduced thiols or further oxidation products (e.g., sulfinic or sulfonic acids), lacking the +89 Da signature. The identification of the modified cysteine was further supported by LC-MS/MS fragmentation analysis, in which the b-and y-ion series confirmed the site of modification **(Fig. 4d** and **Suppl. Fig. 8b)**. An error map showed minimal deviation in fragment ion masses, confirming the confidence of site assignment **(Suppl. Fig. 8c)**.

These results led us to investigate whether *At*PCO4 is also subject to persulfidation *in vivo*. In particular, we asked whether under hypoxia dynamic changes in H₂S could modulate the hypoxic response by inhibiting PCO activity through persulfidation. We introduced a *pPCO4:PCO4:GFP* construct into the *4pco* mutant background, to enable *At*PCO4 immunodetection. Persulfidation levels were quantified as the ratio between persulfidated *At*PCO4 and total *At*PCO4 input using the dimedone switch assay (Aroca et al., 2022). Total protein extracts were split into two fractions, one of which was labelled for persulfidated residues, using the fluorescent probe NBF-Cl/DCP-Bio1, and subsequently purified with streptavidin-bound beads. Both fractions were subsequently probed with an anti-GFP antibody to detect persulfidated *At*PCO4 from the total batch of persulfidated proteins in the extract or total *At*PCO4 input. Seven-day-old seedlings were exposed to hypoxia. Increased *At*PCO4 persulfidation was observed both after 1 and 4 hours of H₂S treatment (**Suppl. Fig. 9**). Interestingly, hypoxia also induced a strong response, particularly at the 4 hours timepoint (**Suppl. Fig. 9**, **Fig. 5a** and **Suppl. Fig. 10a**). In parallel, *At*PCO4 sulfenylation levels in hypoxic seedlings were assessed using the DCP-Bio1 probe, which selectively reacts with sulfenic groups (-SOH), and were quantified relative to the corresponding *At*PCO4 input. Sulfenylation progressively decreased with prolonged hypoxia exposure (**Fig. 5b** and **Suppl. Fig. 10b**), suggesting a redox-based shift in *At*PCO4 from a sulfenylated state under normoxia to a persulfidated state under hypoxia.

**Figure 5.**
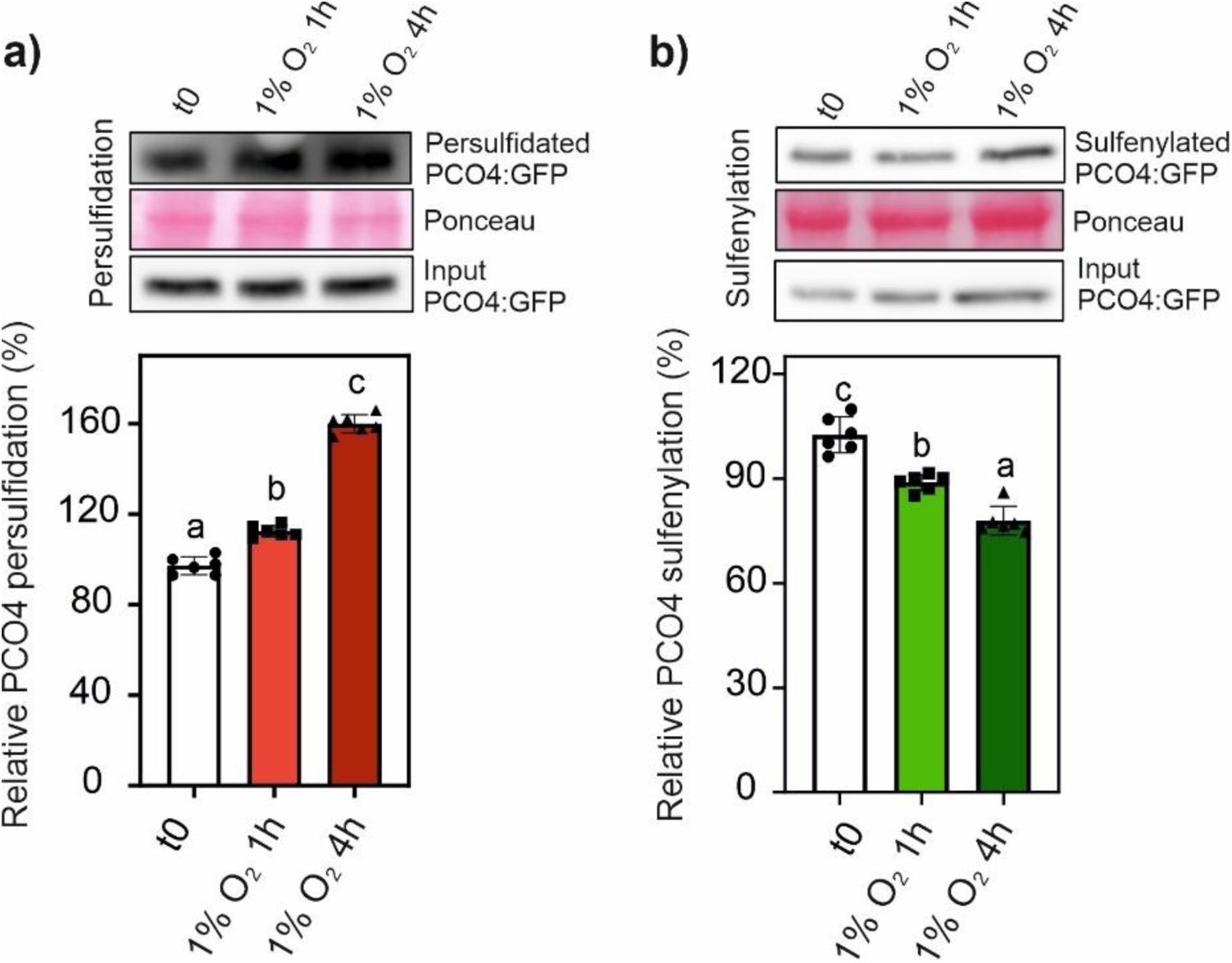
H_2_S-mediated post-translational modification of *At*PCO4 Cys residues. Immunoblot and relative quantification of **a)** persulfidated and **b)** sulfenylated PCO4 in 7-day-old *pPCO4:PCO4:GFP/4pco* seedlings exposed to 1% O₂ for 1 or 4 hours. For a) and b) protein extracts were split into two fractions: one was subjected to specific labeling: (persulfidation: NBF-Cl blocking followed by DCP-Bio1; sulfenylation: DCP-Bio1 labelling of –SOH groups), and the other was used directly as input control. Both fractions were analyzed by immunoblotting with anti-GFP antibodies. Ponceau staining was used as loading control (for further details, see Material and Methods). Data are presented as the ratio between modified and input PCO4 (mean ± SD; n = 6). Different letters indicate statistically significant differences (one-way ANOVA followed by Tukey’s post hoc test). Uncropped blots are shown in **Suppl. Fig. 10**.

### H₂S modulates H_2_O_2_ levels in a dose-dependent and compartment-specific manner

These new insights into S-modification dynamics of PCO4 highlight a potential redox-based shift in PTMs that may influence PCO function under varying oxygen and H₂S conditions. This prompted us to investigate whether H₂S affects H_2_O_2_ levels under normoxic and hypoxic conditions. NaHS supplementation had very mild effects on the induction of transcripts associated with ROS homeostasis, such as *MSD1* and *CAT2* (**Suppl. Fig. 11**), while it stimulated a major response in the transcription factor *ZAT12*, known to be regulated by both oxidative stress and low oxygen (Pucciariello et al., 2012). We used the roGFP2-based biosensors roGFP2-Orp1 and Grx1-roGFP2 to monitor intracellular redox dynamics *in vivo*; these sensors, targeted to the cytosol and mitochondria, allow real-time monitoring of H₂O₂ and glutathione redox potential, respectively (Nietzel et al., 2019).

Experiments were conducted in Col-0 and *erfVII* Arabidopsis seedlings expressing the biosensors under normoxia, hypoxia and reoxygenation, in combination with different concentrations of exogenous H₂S (NaHS).

Under normoxic conditions (21% O₂), H₂S treatment caused a transient increase in cytosolic roGFP2-Orp1 oxidation in a dose-dependent manner, indicating a fast increase in H_2_O_2_ levels (**Fig. 6a**). At lower H_2_S concentrations, sensor oxidation quickly returned to baseline within 30 minutes, suggesting that endogenous antioxidant systems effectively counteract the H₂S-induced oxidative challenge. However, at higher H₂S concentrations (5 mM), a sustained increase in biosensor oxidation was observed. This effect may result from H₂S-induced stimulation of H_2_O_2_ production, or disruption of antioxidant pathways (or both). Also, a direct interaction of H_2_S (and derived polysulfides) (Greiner et al., 2013) with the Cys moieties of the sensor itself cannot be ruled out (Li & Lancaster, 2013).

**Figure 6.**
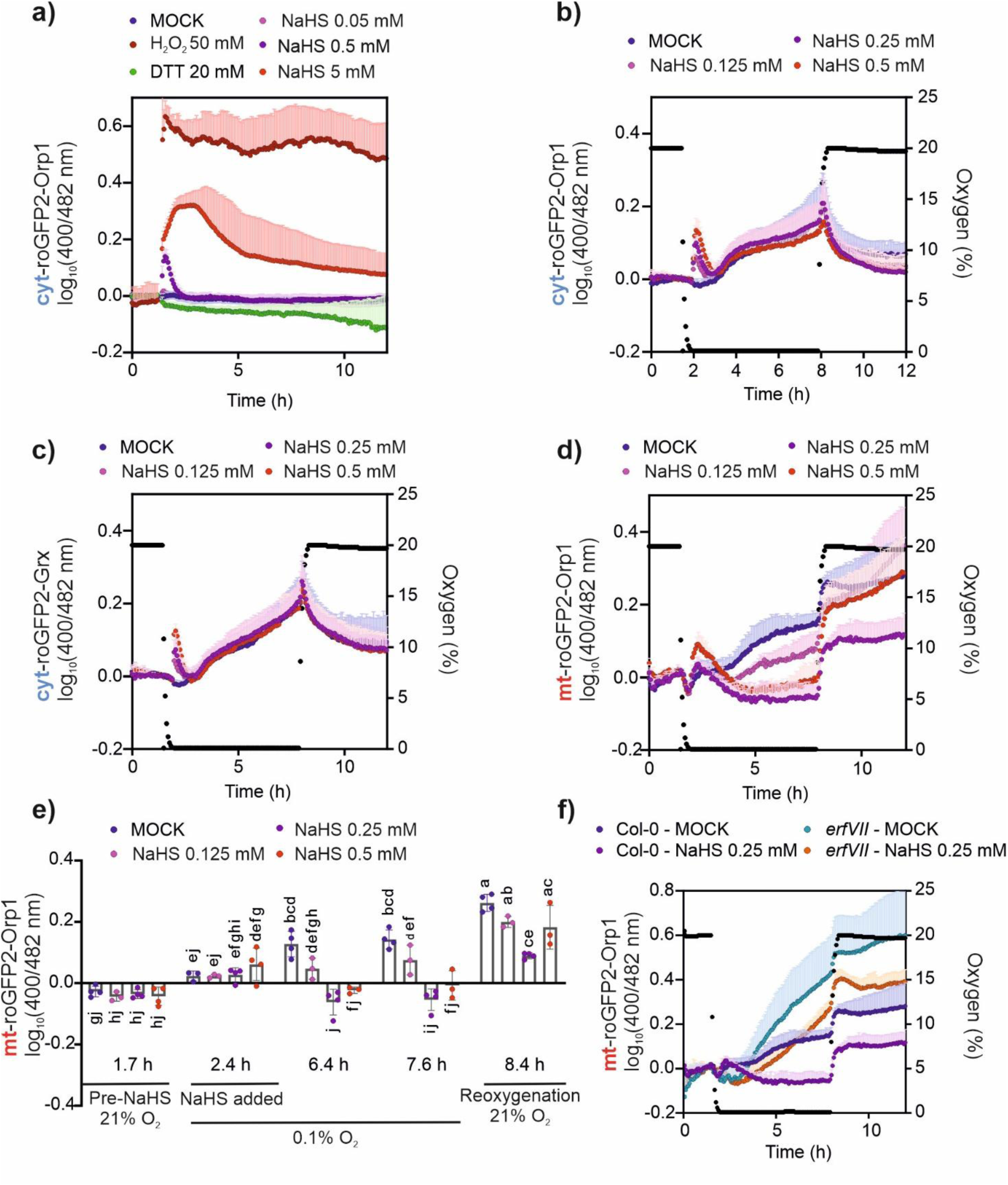
H₂S alters cytosolic and mitochondrial redox states in Arabidopsis under hypoxic conditions. **a)** Time-course measurements of cytosolic redox changes in Arabidopsis seedlings expressing the cytosolic roGFP2-Orp1 sensor treated with varying concentrations of H₂S (NaHS 0.05–5 mM), along with DTT (20 mM, reducing control), H₂O₂ (50 mM, oxidizing control), or mock solution. Data are shown as the log_₁₀_ of the 400/482 nm excitation ratio, indicative of sensor oxidation. **b)** Redox dynamics in cytosolic roGFP2-Orp1-expressing seedlings under mild H₂S stress (0.125– 0.5 mM) in hypoxic conditions (0.1% O₂). Right y-axis shows oxygen concentration (% O₂) over time, indicating the hypoxia treatment and subsequent reoxygenation phase. **c)** Cytosolic redox changes in seedlings expressing Grx1-roGFP2 exposed to 0.125–0.5 mM H₂S under hypoxia. Sensor oxidation is represented as log_₁₀_ (400/482 nm). **d)** Mitochondrial redox changes in seedlings expressing mitochondrial-targeted roGFP2-Orp1 (mt-roGFP2-Orp1) under hypoxia and 0.125–0.5 mM H₂S treatment. Oxygen levels (% O₂) are shown in black right y-axis. **e)** Quantitative analysis of mitochondrial roGFP2-Orp1 oxidation at distinct time points during the hypoxia time course. Seedlings were pre-incubated in normoxia (21% O₂), exposed to H₂S and hypoxia (0.1% O₂), and then reoxygenated. Bars represent means ± SD. Different letters indicate statistically significant differences (ANOVA with Tukey’s post hoc test, *p* < 0.05). **f)** Comparison of mitochondrial matrix redox responses in wild-type (Col-0) (data from panel d) and *erfVII* mutant seedlings expressing mt-roGFP2-Orp1 under hypoxia with or without 0.25 mM H₂S treatment. Oxygen levels (% O₂) are plotted in black on right y-axis.

Under severe hypoxia (0.1% O₂ for 6 h), a similar early H_2_O_2_ increase was detected following H₂S application. However, H₂S did not significantly affect the gradual sensor oxidation associated with hypoxic stress, nor did it modulate the prominent oxidative burst observed during reoxygenation (6 h at 21% O₂) (**Fig. 6b-c**) (Jethva et al., 2023). These effects were consistently observed with both sensors, roGFP2-Orp1 and Grx1-roGFP2-Grx.

Interestingly, H₂S treatment led to less pronounced oxidation of sensors expressed in the mitochondrial matrix under hypoxia, suggesting that H₂S may attenuate mitochondrial H₂O₂ production or enhance specific antioxidant responses, or both (**Fig. 6d, e**). This is consistent with prior studies describing H₂S as a mitochondrial electron donor and as an inhibitor of cytochrome *c* oxidase, both of which could influence mitochondrial ROS generation (Cooper & Brown, 2008; Fu et al., 2012.; Pedroletti et al., 2023; Szabo, 2010)

Despite this, the reoxygenation (6 h at 21% O₂)-induced burst of oxidation in the matrix remained unaffected by H₂S. To determine whether the ERFVII transcription factors influence this H₂S-mediated redox response in mitochondria, we compared the redox sensor dynamics in Col-0 and *erfVII* mutant backgrounds using mitochondrial probes. Even though the progressive oxidation during hypoxia was strongly increased in the *erfVII* mutant as compared to the wild-type, the responses to H₂S treatment were similar between genotypes (**Fig. 6f**), suggesting that this apparent ability of H₂S to counteract sensor oxidation in the matrix is independent of ERFVII signalling pathways.

Collectively, these results indicate that H₂S plays a context- and concentration-dependent role in cellular redox regulation. While H₂S can transiently increase H_2_O_2_ under normoxic and hypoxic conditions, it may also serve a protective role in mitochondria by dampening H₂O₂ production under hypoxia, a phenomenon that should be further investigated. These findings provide mechanistic support for a model in which H₂S modulates redox status through both pro-oxidant and antioxidant mechanisms, with implications for PCO4 function and hypoxia signalling.

### Decreased H₂S levels are associated with a weaker hypoxia responses and lower tolerance to low-oxygen conditions

The increase in PCO4 persulfidation in seedlings exposed to hypoxia (**Fig. 5a**) hints at a possible impact of H₂S on the establishment of the hypoxic response to low oxygen deprivation. Free H₂S quantification in Col-0 seedlings exposed to hypoxia showed a decrease after both 1 hour and 4 hours treatment (**Fig. 7a**). This may be due to its utilization for the formation of persulfide groups on Cys residues of proteins, which may help protect them from irreversible oxidation caused by ROS accumulation under hypoxic condition (Pucciariello et al., 2012). We adopted a genetic strategy to verify whether the physiological H₂S levels present in seedlings under hypoxia may contribute to determine the dynamics of hypoxic responses. We exposed the *des1* mutant, characterized by a 30% reduction in free H_2_S (Álvarez et al., 2012), to short-term hypoxia (1% O_2_), followed by analysis of HRG expression. In the mutant, the early induction of most transcripts tested was significantly lower than in Col-0 (**Fig. 7b**). All markers but *PCO1* exhibited lower expression in *des1* after 1 hour of hypoxia exposure, whereas *HRA1* decrease was already visible after 30 minutes (**Fig. 7b**). We also evaluated impact of *des1* under submergence, a condition that favours the entrapment of gaseous signals like H₂S. Seedlings were completely covered with water to simulate a dark flooding condition in the well. Also in this case, a less pronounced hypoxic response was observed in the *des1* mutant compared to Col-0 (**Fig. 7c**). No differences were seen after 30 minutes, but all genes except *ADH1* showed lower expression in *des1* 1 hour into the treatment (**Fig. 7c)**.

**Figure 7.**
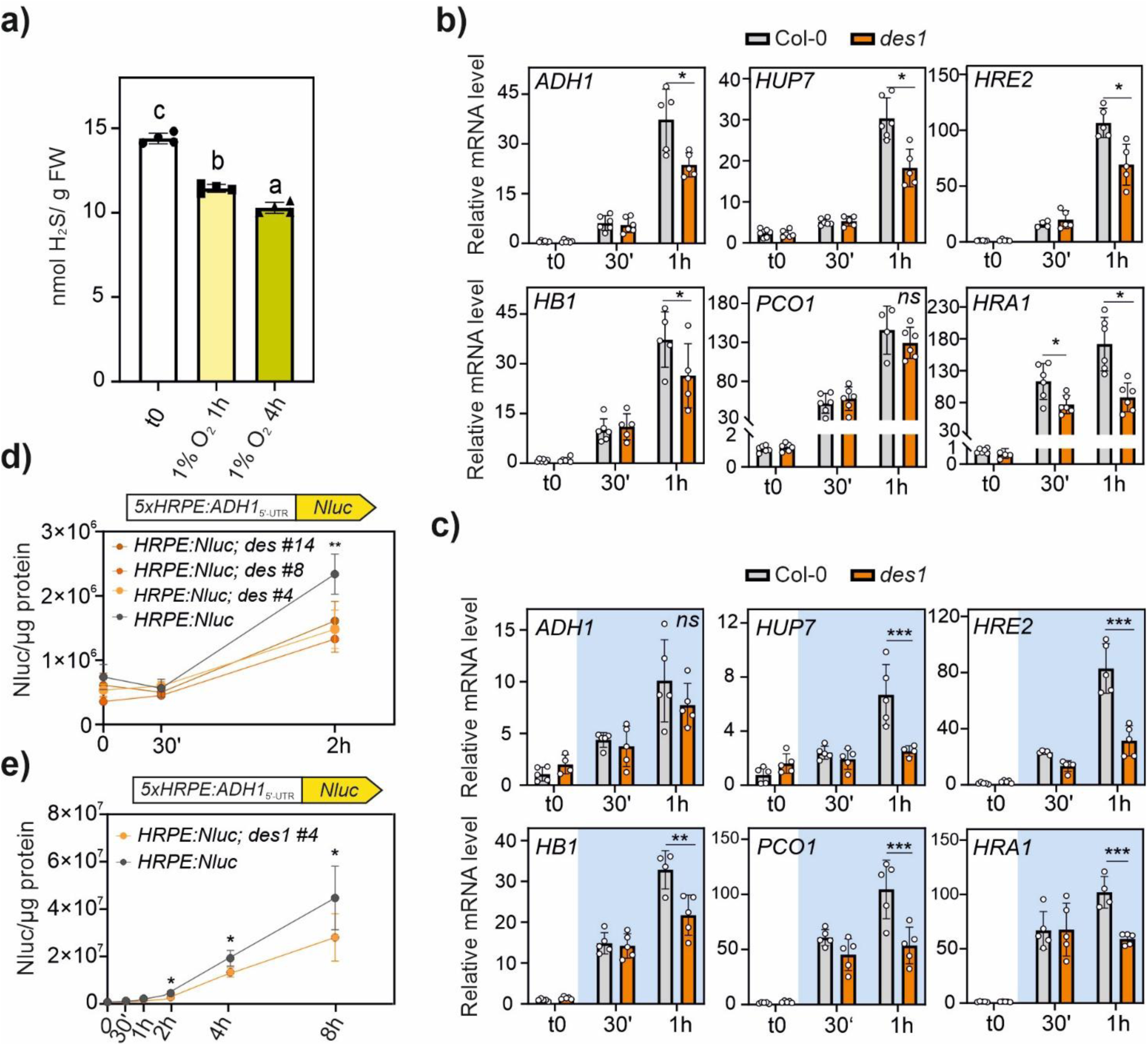
H_2_S involvement under low oxygen. **a)** Relative quantification of free H_2_S on fresh weight (FW) in 7-days old Col-0 seedlings exposed to dark hypoxia (1% O_2_ v/v) for 1 or 4 hours. Data are mean ± SD (n=4). Different letters denote significant differences (one-way ANOVA followed by Tukey’s post hoc test). **b)** HRG expression in 7-days old Col-0 and *des1* seedlings exposed to dark hypoxia (1% O_2_ v/v) for 30 minutes or 1 hour. In control condition seedlings were treated in the dark at atmospheric oxygen concentration (21% O_2_ v/v). Data are mean ± SD (n=5). **c)** HRG expression after 30 minutes or 1 hour submergence. Controls samples (t0) were treated in the dark under 21% O_2_. **d)** Luciferase activity in three independent *HRPE*-based reporter lines in *des1* background. *HRPE:Nluc* seedlings in Col-0 or *des1* mutant background were grown for seven days on vertical plates and treated with dark hypoxia (1% O_2_ v/v) for 30 minutes or 2 hours. **e)** Extended response to hypoxia in 7-days old *HRPE:Nluc #4* seedlings in *des1* or Col-0 background. Nanoluciferase signals were normalised to the total soluble protein (Nluc/µg total protein); data are mean ± SD (n=4). Asterisk indicates statistically significant differences between genotypes after Student’s t-test (*p<0.05; **p<0.01; *** p<0.001).

We further characterized the dynamics of the hypoxic responses in the mutant by introducing the *HRPE*:*Nluc* reporter in the *des1* background. Three independent lines showed a comparable signal when exposed to low oxygen, with a lower signal after 2 hours of hypoxia compared to the Col-0 background (**Fig. 7d**). We selected the *HRPE:Nluc #4* to monitor the response under prolonged hypoxia. The signal showed a lower intensity after 2, 4, and 8 hours of exposure to 1% of O_2_ than in the wild-type background (**Fig. 7e**).

Given the accumulating evidence suggesting a role for H₂S in mediating the hypoxic response during oxygen deprivation, we tested the consequences of H_2_S biosynthesis impairment on submergence tolerance. The *des1* mutant showed lower stress tolerance as compared to the wild-type after 60 hours dark submergence (**Fig. 8a, Suppl. Fig. 12)**. No significant differences were instead observed following 72 hours of submergence, which represented a lethal treatment for both genotypes. The survival rate after 1 week of recovery from 60 hours of submergence was 93% for Col-0, whereas it was only 26% for the *des1* mutant (**Fig. 8b**). Additionally, under these conditions, the mutant showed a significant decrease in Plant Leaf Area (PLA) compared to the wild-type, a difference that was not observed either in the controls or after 72 hours of submergence (**Fig. 8c**). Together, these findings suggest a role for H_2_S in modulating the hypoxic response, emphasizing how appropriate levels of H_2_S contribute to the initiation and maintenance of an adequate response during oxygen deprivation.

**Figure 8.**
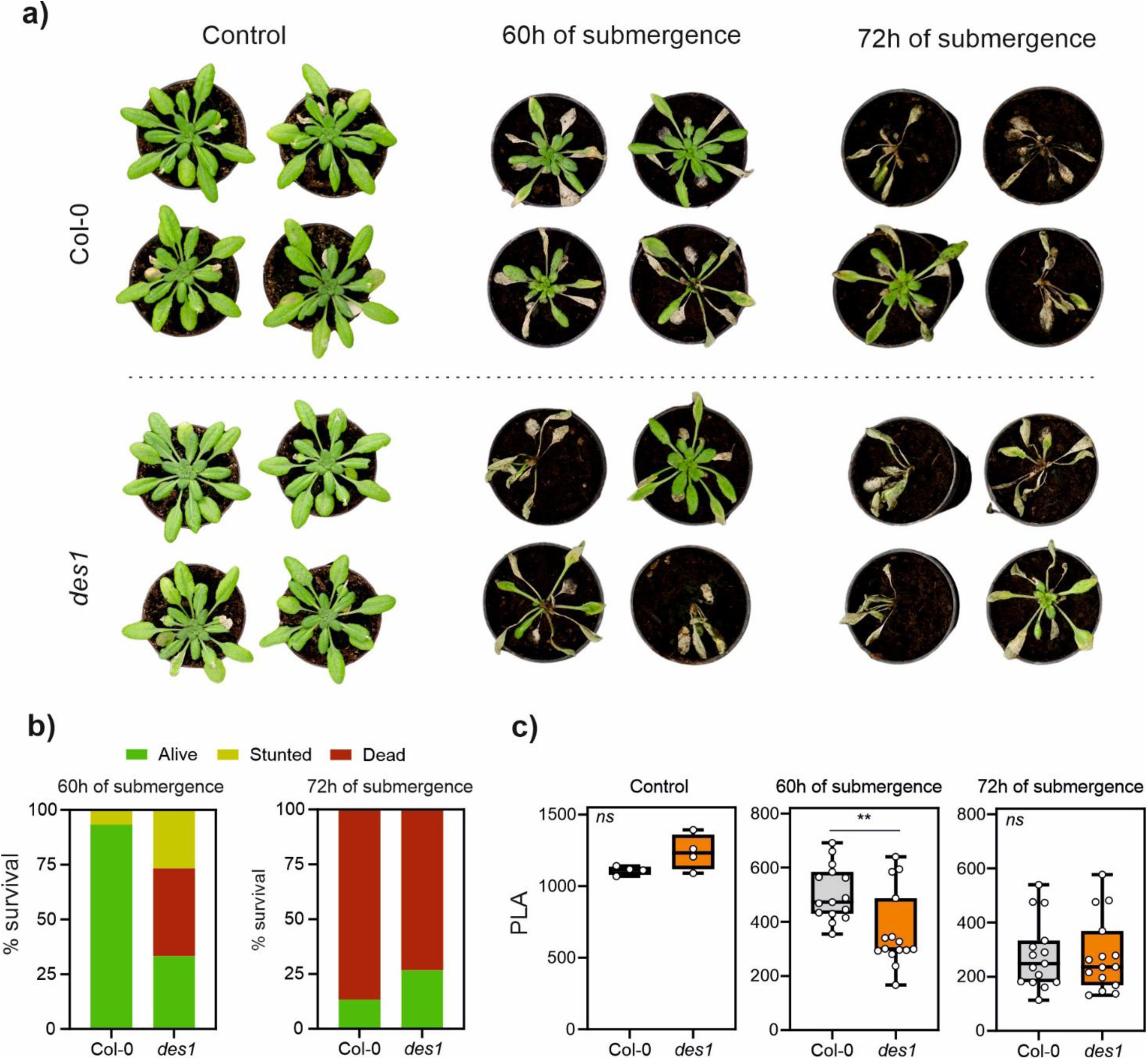
Effect of submergence on Col-0 and *des1* plants. **a)** Images of representative Col-0 and *des1* plants after 1 week of recovery from dark submergence. The control plants were treated under dark conditions for 60h. **b)** Survival rate of Col-0 and *des1* after 1 week of recovery from submergence. Categories: Alive = all or most old leaves alive and new leaves produced; stunted = new leaves produced but most or all old leaves dead; dead = all or most old leaves dead, no or very few new leaves (n=4 for control, n=15 for submergence). **c)** Plants Leaf Area (PLA) in the Col-0 and *des1* after 1 week from the submergence. Data are mean ± SD (n=4 for the control, n = 15 for the treated plants). The asterisks indicate a statistical difference after the student’s t-test (*0.01≤p<0.05; **0.001≤p<0.01; ***p<0.001).

## Discussion

Once considered as a toxic gas, H₂S is now recognized for its regulatory role in several physiological processes and responses to both biotic and abiotic stresses, such as salinity, drought and waterlogging (Chen et al., 2015; Jurado-Flores et al., 2023; Y. Li et al., 2021). Although several studies have reported that exogenous H₂S can mimic aspects of the hypoxic response and improve survival rate of *A. thaliana* plants exposed to submergence (Xiao et al., 2020; Li et al., 2021; Zhou et al., 2021), its role as an endogenous modulator of hypoxic signalling remains poorly understood. Here, we describe PCO enzyme persulfidation as a molecular mechanism connecting H₂S with the modulation of the hypoxic responses in *A. thaliana*.

We first investigated whether alterations in sulfate assimilation, a primary route of endogenous H₂S production (Aroca et al., 2018), could influence the hypoxic response. Bielecka et al. (2015) have previously reported HRG up-regulation in *A*. *thaliana* seedlings grown under S-deficient conditions early after S re-supplementation (**Suppl. Table 1**). Notably, no significant changes in metabolite levels were detected within the first hour of nutritional shift, suggesting that signalling events, rather than metabolic regulation, should have caused HRG induction. Our results indicate that restoration of S following S starvation caused induction of the hypoxic genes, under aerobic conditions, in an ERFVII-dependent manner (**Fig. 1b-d**). Having ruled out the occurrence of local hypoxia due to increased O_2_ consumption (**Fig. 1e, Suppl. Fig. 1**), we put forward that the observed hypoxia-like responses might be caused by a short-term direct effect of S re-integration on the O_2_-sensing machinery.

The involvement of H₂S in this phenomenon was confirmed through the application of sodium hydrosulfide (NaHS), as H₂S source. A minimum concentration of 100 µM NaHS, a non-toxic dose commonly applied for short term treatments in Arabidopsis (Zhou et al., 2025) was sufficient to stabilize RAP2.12 and promote HRPE-driven reporter expression under normoxic conditions (**Fig. 2a-c**). The induction of hypoxic genes was weak at 100 µM, but became progressively more pronounced at higher concentrations, highlighting the transient and dose-dependent effect of the exogenous treatment (**Fig. 2d**). High concentrations of NaHS (e.g. 500 µM) also triggered the expression of oxidative stress marker genes (**Suppl. Fig. 11**), suggesting a ROS burst. Although H₂S is well established as a ROS scavenger (García-Calderón et al., 2023; Jaiswal et al., 2024; Wang et al., 2022), some studies indicate that it can also trigger indirect ROS production when supplied at high doses (Shen et al., 2020; Zhang et al., 2017). Consistently, NaHS supplementation resulted in detectable cellular redox changes, as evidenced by the roGFP2-derived biosensors (**Fig. 6**), under both hypoxic and normoxic conditions. Recently, the inhibitory effect of H₂O₂ on PCO activity has been demonstrated (Akter et al., 2024). Therefore, we cannot rule out that the indirect mediation of ROS might concur to the promotion of hypoxia-like responses by H₂S. However, HRG expression patterns were maintained all over the range of H₂S concentration (**Suppl. Fig. 4-6**), both at low doses, that are likely to be associated with ROS scavenging, and at higher doses, that might trigger oxidative stress. This suggests that the direct effect of H₂S might be predominant over any indirect effects mediated by ROS.

Persulfidation of susceptible Cys residues are considered the predominant mechanism mediating H_2_S-dependent signalling, through changes in target protein biochemical activity, interactions or structure (Shen et al., 2020; Wang et al., 2023). Rapid induction of HRG expression can follow from the inhibition of PCO enzymes (Lavilla-Puerta et al., 2023), within minutes of hypoxia (Lavilla-Puerta et al., 2025). We thus hypothesized that fast stimulation of hypoxic responses by H₂S (**Fig. 2c-d**) might be linked to the persulfidation of PCOs. In a previous proteomic study, Aroca et al., (2017) have identified PCO3 as a target of both S-nitrosylation and persulfidation, highlighting the potential for redox regulation within this enzyme family. Compatibly, we observed persulfidation on a recombinant *At*PCO4 *in vitro* after H₂S incubation (**Fig. 4**), leading to the inhibition of its Cys oxidation activity. Although the concentrations of *At*PCO4 and H₂S are likely to be much lower *in vivo* than in our experiments with recombinant enzyme, we nevertheless observed an inhibitory effect that was proportional to H₂S dose and exposure time (**Fig. 3a-c**). The inhibitory effect was present on all Arabidopsis PCO isoforms and partially reversible by reducing agents, supporting the model of a redox-dependent inhibition of PCO enzymes through persulfidation (and likely oxidised derivatives thereof) (**Fig. 3d-f**). Through LC-MS/MS, we identified Cys172 as a *At*PCO4 persulfidation site. While we cannot rule out that other Cys residues may also be persulfidated, this conserved residue has been shown to play a role in enzyme interaction with O₂ and catalysis (Dirr et al., 2025). The *At*PCO4-2 C173A variant (corresponding to *At*PCO4-1 Cys172) exhibited lower catalytic efficiency of dioxygenation *in vitro* than the wild-type version, without affecting substrate binding, and, when expressed in Arabidopsis, conferred higher tolerance to submergence (Dirr et al., 2025). Our analysis revealed that Cys172 can exist in several oxidized forms (**Fig. 4c**), in line with its proximity to the active site and confirming its interaction with O_2_. The fact that Cys172 can undergo irreversible oxidation (**Fig. 4**), but is also susceptible to persulfidation, suggests a protective role of H₂S in maintaining *At*PCOs functionality under oxidative conditions (Vignane & Filipovic, 2023).

We also observed *in vivo* that persulfidation levels of *At*PCO4 increased in seedlings exposed to NaHS and are further elevated under hypoxic conditions (**Suppl. Fig. 9**), suggesting a direct involvement of H₂S as physiological messenger in the modulation of hypoxic responses. Enhanced persulfidation was accompanied by a decrease in sulfenylation levels (**Fig. 5a-b**), supporting the idea of a finely tuned redox regulation of *At*PCOs according to oxygen availability. Although counterintuitive, ROS accumulation is among the common consequences of hypoxia and has been detected as early as 30 minutes into oxygen deprivation (Gonzali et al., 2015). The ROS-scavenging properties of H₂S, which under hypoxia are supposed to help mitigate oxidative stress caused by ROS excessive production, seem to be reflected by the observed decrease of H₂S levels under hypoxia (**Fig. 7a**). Rapidly synthesized H₂S might thus have been consumed to increase the persulfidation levels in hypoxia (**Fig. 5a**), resulting in lower steady-state levels of H₂S after 1- or 4-hours hypoxia. Previous studies have shown H₂S accumulation under low-oxygen conditions, particularly during submergence or anoxia (Yang et al., 2021; Zhou et al., 2021). However, these conditions may induce effects not taking place in our experimental setting of gaseous hypoxia (**Fig. 5a**), such as the local entrapment of gaseous molecules due to water coverage. A comparative analysis of H₂S dynamics under different oxygen-deficient conditions could help clarify the discrepancy observed in current data.

Use of the *des1* mutant (Álvarez et al., 2010) impaired in Cys desulfhydration capacity in the cytosol, underscored the importance of H₂S as a physiological messenger during the establishment of the anaerobic response. Lowering of the cytosolic H₂S levels impacted on HRG induction under low oxygen (**Fig. 7b-e**), compatible with the (partial) preservation of PCO activity in comparison with wild-type seedlings. Furthermore, *des1* impaired tolerance to submergence (**Fig. 8**) suggests a broader role for H₂S in low-oxygen stress, beyond the inhibition of PCOs to enable full activation of transcription following hypoxia. H_2_S impact under low oxygen might extend to safeguard key proteins from irreversible oxidation occurring in the post-submergence phase of reoxygenation, thereby helping plant recovery after stress. Quantitative analysis of the persulfidated proteome (persulfidome) under submergence would provide valuable insights into this hypothesis.

Here, we provided the first direct evidence for an endogenous role for H₂S to set up the hypoxic response in plants and highlights its contribution to the complex biochemical and molecular networks for survival under submergence stress.

## Methods

### Plant material

The Columbia-0 ecotype (Col-0) was used as the wild-type background in all experiments. The *des1* mutant (SALK_103855) was obtained from NASC (National Arabidopsis Stock Center). The quintuple *erf-VII* mutant was previously described in Abbas et al. (2015). The transgenic lines *35S:RAP2.12_1-28_:Fluc* (Licausi et al., 2011), *35S:RAP2.3^3xHA^* (Gibbs et al., 2014) and the *HRPE:Nluc* (Akter et al., 2024) were previously described. The *HRPE:Nluc*/*des1* line was obtained by crossing *des1* with the *HRPE:Nluc* line. The homozygous seeds were selected on ½ MS supplemented with 10 g l^−1^ of sucrose, 9 g l^−1^ of agar and glufosinate ammonium 1mM. F_3_ seeds were collected and used for the experiments.

Wild-type *Arabidopsis thaliana* lines stably expressing the cytosolic and nuclear-localized roGFP2-Orp1 biosensor were previously described (Nietzel et al., 2019). To generate sensor lines in the *erfVII* background, Agrobacterium tumefaciens-mediated floral dip transformation was performed using a pH2GW7:mt-roGFP2-Orp1 construct. Transgenic lines were selected on hygromycin B-containing media and screened for sensor fluorescence.

For the generation of the PCO4:GFP construct, a 1519-nt fragment was synthesized by Integrated DNA Technologies (IDT), including the *attB1* and *attB2* sequences upstream and downstream of the *PCO4:GFP* sequence, respectively. The coding sequence corresponding to the PCO4-1 version was adopted. The fragment was first cloned into pDONR207 and subsequently into pH7pPCO4GW (White et al., 2020) via BP and LR reactions, respectively, following the manufacturer’s instruction. For sequence details see **Supplemental Table 3.** Agrobacterium-mediated transformation using the floral dip method (Clough & Bent, 1998) was employed to generate a stable *pPCO4:PCO4:GFP* transgenic line in the *4pco* background (Masson et al., 2019). T_0_ seeds were selected for hygromycin resistance on solid MS medium and subsequently transplanted into soil. Integration of the construct was confirmed by PCR using GoTaq® DNA polymerase (Promega). Experiments were conducted using seeds from the T_3_ generation.

### Growth conditions

For axenic experiments, seeds were sterilized with ethanol 70% (v/v) for 1 minute, incubated in 10% (v/v) sodium hypochlorite (NaClO) for 10 minutes, followed by 5 times washes with sterile water. Seeds were incubated in the dark for 48 h at 4°C before the growing. For the nutrient reintegration experiments, seedlings were grown aerobically in 6-well plates for 5 days in 2.5 mL of deficiency liquid medium and then transferred into 2.5 mL of complete liquid media. The media compositions are described in **Supplemental Table 2**. Experiments in Figure 2 and Figure 4b, 4c were conducted on 7-days old seedlings grown for 6 days on ½ MS (Duchefa) supplemented with 10 g l^−1^ di sucrose and 9 g l^−1^ agar (Duchefa) and transferred for one day in 6-well plates with 2.5 mL of ½ MS (Duchefa) with the addition of 10 g L^−1^ of sucrose. Soil experiments were carried out on soil:vermiculite 3:1 ratio mixture. Plants were grown under atmospheric conditions with 23°C day/19°C night temperature and a 12 h light period with a light intensity of 120 μmol photons m^−2^s^−1^.

### Low oxygen treatments

For hypoxia treatments, seedlings were grown on vertical plates for 7 days and exposed with 1% O_2_/N_2_ (v/v) atmosphere recreated in a sealed glovebox (Coy), or normoxic atmosphere, for the indicated duration, in the dark. In partial submergence experiments, the seedlings were grown for 6 days on vertical plates and transferred for one day into 6-well plates. Subsequently, the media was removed, and the seedlings were covered with 5 mL MilliQ water and exposed to 1% O_2_/N_2_ (v/v) in darkness for the specified duration. Seedlings in control conditions were exposed to dark and atmospheric oxygen without the removal of the liquid media. All the treatments were performed using the Gloveless Anaerobic chamber (COY Laboratory Products). The submergence was performed in 21-days old plants grown in soil. After 60h or 72h of dark submergence the photoperiodic period was restored (23°C day/19°C night temperature and a 12h light period with a light intensity of 120 μmol photons m^−2^ s^−1^). Control plants were exposed to dark conditions for 60h. Survival rate was determined using the following parameters: “Alive”: all or most old leaves alive, new leaves produced; “Stunted”: new leaves produced, but most or all old leaves dead; “Dead”: all or most old leaves dead, absence of new leaves. Before the submergence and after 1 week of recovery the Plant Leaf Surfaces (PLA) was calculated using a LabScanalyzer digital phenotyping machine (LemnaTec GmbH, Aachen, Germany) equipped with a Manta G-1236 camera and a Kowa LM12XC lens.

### Chemical treatments

H_2_S treatments were induced by supplementation of NaHS (Sigma-Aldrich) dissolved in MilliQ water, freshly prepared. Pyruvate treatments were performed by supplementation of 8 mM pyruvate (Sigma-Aldrich) dissolved in MilliQ water.

### O_2_ consumption measurements

O_2_ consumption over time was assessed in seedlings placed inside cuvettes (Pyroscience) containing 2 mL of sterile water, kept stirring with a magnet. The variation of O_2_ concentration was measured with a phosphorescent sensor (Fiber-Optic Oxygen Meter 2 channels FSO2-C2, Pyroscience), which detected the data through an O_2_ sensitive spot attached on the internal surface of the cuvette. The collected data were recorded and analyzed using the software PyroDataInspector. To compare O_2_ consumption between different samples, the parameter Oxygen Consumption Indicator (OCI) was introduced, defined as the time required for dissolved O_2_ to decrease from 85% to 70% saturation (in water initially equilibrated with a 21% O_2_ atmosphere), normalized with respect to the fresh weight.

### Biosensing of intracellular H_2_O_2_ dynamics in living Arabidopsis tissue

Mt-roGFP-Orp1 sensor lines in the *erfVII* background were used in both T_2_ (segregating for biosensor) and T_3_ (homozygous) generations, depending on the experiment: T_2_ lines for leaf disc assays and T_3_ for seedling experiments. Wild-type sensor lines were homozygous for the insertion. To account for positional effects, two independent sensor lines per genotype were analyzed, and replicates were pooled for final data analysis. Leaf discs (from 5-week-old plants) and 7-day-old seedlings were submerged in 200 µL standard assay buffer (10 mM MES, pH 5.8 adjusted with KOH, 10 mM MgCl₂, 10 mM CaCl₂, 5 mM KCl) in a 96-well plate. Each well received a single leaf disc (abaxial side up) or 5–6 seedlings. Ratiometric fluorescence measurements were performed using a ClarioStar Plus microplate reader (BMG Labtech) in top-optic mode with 30 excitation flashes and 3 mm averaging diameter (orbital for discs, spiral for seedlings). Excitation/emission settings were: Ex1: 400 ± 10 nm, Ex2: 482 ± 16 nm; Dichroic mirror: LP504; Em: 520 ± 10 nm. Data were collected every 243 seconds. Oxygen concentration was controlled using an Atmospheric Control Unit (BMG Labtech) by targeted N₂ influx. Separate experimental repetitions were used for each O₂ treatment. Background correction was performed using wildtype Col-0 samples and background was subtracted before calculating 400/482 nm ratios. Ratio data were log₁₀-transformed for statistical normalization.

### RNA extraction and gene expression analysis

RNA extraction was performed as previously described (Kosmacz et al., 2015). The RNA quality and quantity were checked by gel electrophoresis on 1% (w/v) agarose and by spectrophotometric analysis. Maxima First-Strand complementary DNA (cDNA) Synthesis Kit (Thermo Fischer Scientific) was used for the reverse transcription, according to the manufacturer’s recommendation. ABI Prism 7300 sequence detection system (Applied Biosystems) was used for the RT-qPCR using 12.5 ng cDNA with PowerUp SYBR® Green Master Mix (Thermo Fisher Scientific). The relative gene expression was calculated using UBQ10 (AT4G05320) as housekeeping gene through the ΔΔCt method (Livak & Schmittgen, 2001). Primer sequences are specified in **Supplemental Table 3**.

### Luciferase assays

Seedlings were ground in liquid nitrogen and the total protein extracted in passive lysis buffer (Promega). ONE-Glo Luciferase Assay kit (Promega) was used to quantify the Firefly luciferase activity and the NanoLuc luciferase enzyme was measured with the Nano-Glo® Luciferase Assay System (Promega). Luciferase signals were normalized on the total protein concentration quantified using Bradford assay (Bradford, 1976).

### Persulfidation and sulfenylation detection

Protein extracts were obtained from plant material in 1x phosphate-buffered saline (PBS) pH 7.4, supplemented with 1 mM EDTA, 2% w/v SDS, and 1x protease inhibitor cocktail (Thermo). The total amount of protein was determined using the DC Protein Assay (Bio-Rad). The protein extract was divided into two batches, one of which was used for immunoblot analysis (input extract) and the second was subjected to the dimedone switch method followed by immunoblot analysis (persulfidated and sulfenylated extract). The persulfidation and sulfenylation of PCO was detected after treatment following the dimedone switch method previously described by Aroca et al., (2022). Briefly, for persulfidation analysis, NBF-Cl (Merck) was employed to derivatize all cysteine residues and amino groups, and then a labelling step with DCP-Bio1 (Merck) was performed for tagging the persulfidated adducts. The labelled proteins were purified by incubation with Sera-Mag Magnetic Streptavidin beads (Cytiva) overnight at 4 °C and subsequent elution from the beads with 2.25 M NH_4_OH and 2% SDS at 95°C for 10 minutes. Eluted proteins were then precipitated with acetone/TCA and resuspended in 1x PBS and 2% SDS. Proteins of the input and the persulfidated extract were separated in an SDS–PAGE 10% (w/v) polyacrylamide gel and transferred to a nitrocellulose membrane. PCO:GFP was detected with the antibodies anti-GFP (Bioscience, dilution 1:1000) in PBS containing 0.1% Tween 20 (Sigma-Aldrich) and 5% milk powder. The ECL Select Western blotting Detection Reaction (GE Healthcare) was used to detect proteins with horseradish peroxidase-conjugated anti-rabbit secondary antibodies. For a protein loading control, the membrane before immunodetection was stained with Ponceau S (Sigma-Aldrich) to detect all protein bands. Sulfenylated proteins were detected *via* DCP-Bio1 labelling and visualized using Alexa Fluor™ 488-conjugated streptavidin, as previously described (Zivanovic et al., 2019). Sulfenylation levels were quantified using β-tubulin, detected by western blotting (As10680; Agrisera, Vännäs, Sweden; dilution 1:5000), as an internal loading control.

### H_2_S quantification

100 mg of leaf tissue were homogenised in liquid nitrogen and metabolites were extracted in homogenous mixture of Tris-HCl buffer (100 mM, pH 8.5; EDTA 1 mM), with shaking for 30 min at 4 ◦C. Samples were then sonicated for 5 min in ice-bath and centrifuged for 15 min at 12,500× g at 4 ◦C; 100 µL of supernatants were derivatised with 25 µL of 15 mM monobromobimane (MBB, Merck) solution for 30 min at room temperature, and stopped by adding 25 µL of 5% formic acid. The mixture was subjected to centrifugation at 800× g for 10 min, and 1 µL of the supernatant was injected into the LC–MS/MS system for analysis. A calibration curve for NaHS concentrations was established ranging from 2.5 µM to 100 µM, and the H_2_S concentration in the samples was determined using this standard curve. The results are presented mean values ± SD of three different biological replicates.

### Recombinant protein production

Recombinant PCO proteins from *A*. *thaliana* were expressed and purified following previously established protocols (White et al., 2018). His6-tag affinity purification was performed, after which the His6-tag was cleaved using TEV protease and subsequently removed with a HisTrap HP column (GE Healthcare). The proteins underwent further purification using a HiLoad 26/600 Superdex 75 prep-grade size exclusion column (GE Healthcare) equilibrated with 50 mM Tris (pH 7.5) and 0.4 M NaCl. The purity of the isolated proteins was confirmed by SDS-PAGE.

### *In vitro* H_2_S oxidation assay of PCOs

For the *in vitro* H₂S oxidation assay, 10 μM recombinant PCO (*At*PCO1-5) was incubated with defined concentrations of H₂S, H₂O₂, or an equivalent volume of H₂O at 4 °C for specified time duration. Subsequently, 200 μM RAP2_2-15_ peptide (CGGAIISDFIPPPR, purchased from GL Biochem, China) was reacted with 1 μM PCO (treated or untreated with H₂S/H₂O₂) at 25°C for the desired duration in 50 mM HEPES buffer (pH 7.4), hereafter referred to as reaction buffer. The reaction was halted by quenching 5 μL of the sample in 45 μL of 5% formic acid, and peptide masses were analyzed using an Agilent RapidFire RF360 sampling robot coupled to an Agilent 6530 Accurate-Mass Q-ToF mass spectrometer operating in positive electrospray mode. The distribution of reaction products was determined based on the relative integrated areas of corresponding peaks, and spectra were visualized using Qualitative Analysis (Version B.07.00). Agilent RapidFire Integrator (Version 4.3.0.17235) was used to compute the integrated peak areas.

A related experiment examined the protective effects of GSH or TCEP against H₂S-mediated inhibition of PCO activity. In this experiment, 2 μM PCO was treated with or without 5 mM H₂S for 30 minutes on ice, followed by 10 mM GSH/TCEP treatment (30 minutes on ice). Control samples, which were not exposed to H₂S, were simultaneously treated with 10 mM GSH. Following treatment, 1 μM *At*PCO was reacted with 100 μM *At*RAP2_2-15_ at 25°C for defined time periods in the reaction buffer. Reactions were quenched with 5% formic acid, and two replicates were performed for each condition.

### Detection of Cys modification by H_2_S *in vitro*

To detect Cys oxidative modifications in AtPCO4, BioDiaAlk, a biotinylated form of DiaAlk which is a chemical probe specific to sulfinic acid formation, was utilized. 100 μM of recombinant AtPCO4 enzyme was incubated with H₂S, H₂O₂, or an equivalent volume of H₂O in reaction buffer for 1 hour at 25°C. Excess H₂S/H₂O₂ was removed using a Micro Bio-Spin P-6 chromatography column (Bio-Rad) equilibrated with reaction buffer. AtPCO4 samples were then incubated with 1 mM BioDiaAlk in the dark for 1 hour at 25°C, followed by reduction with 10 mM DTT for 1 hour at 25°C. Proteins were separated using SDS-PAGE, transferred onto a polyvinylidene difluoride (PVDF) membrane, and subjected to streptavidin-HRP blotting (1:1000 dilution) or anti-His-HRP blotting (1:10,000 dilution). The protein signals were visualized using chemiluminescence (ECL Plus, Pierce).

### Trypsin digest and LC-MS/MS to detect AtPCO4 modification sites

To identify modification sites on AtPCO4, 15 μg recombinant enzyme was incubated with H_2_S or H_2_O_2_ or equal volume of H_2_O in reaction buffer for 1 h at 25°C, followed by 100 mM iodoacetamide (IAM) treatment to block free thiols for 1h in dark at 25°C. After removing excess H_2_S/H_2_O_2_/IAM by spin column, in-solution trypsin digestion was performed by adding trypsin in a 1:50 (w/w) ratio overnight at 37 °C, followed by 85 mM DTT reduction for 1 h at RT to break the persulfide bridge. Reduced peptides were purified by C18 ZipTip column and resulting tryptic peptides were resuspended in 40 μL of Milli-Q water with 2 % acetonitrile and 0.1 % formic acid. 2 µL resuspended peptides were analysed on a NanoAcquity-UPLC system (Waters) connected to an Orbitrap Elite mass spectrometer (Thermo Fischer Scientific) possessing an EASY-Spray nano-electrospray ion source (Thermo Fischer Scientific). The peptides were trapped on an in-house packed guard column (75 μm i.d. x 20 mm, Acclaim PepMap C18, 3μm, 100 Å) using solvent A (0.1 % Formic Acid in water) at a pressure of 140 bar. The peptides were separated on an EASY-spray Acclaim PepMap® analytical column (75 μm i.d. × 50 mm, RSLC C18, 3 μm, 100 Å) using a linear gradient (length: 100 minutes, 3 % to 60 % solvent B (0.1 % formic acid in acetonitrile), flow rate: 300 nL/min). The separated peptides were electrosprayed directly into the mass spectrometer operating in a data-dependent mode using a CID-based method. Full scan MS spectra (scan range 350-1500 m/z, resolution 120000, AGC target 1e6, maximum injection time 250 ms) and subsequent CID MS/MS spectra (AGC target 5e4, maximum injection time 100 ms) of 10 most intense peaks were acquired in the Ion Trap. CID fragmentation was performed at 35 % of normalised collision energy, and the signal intensity threshold was kept at 500 counts. The CID method used performs beam-type CID fragmentation of the peptides. Data analysis was performed with Peaks 8.5. The raw MS file was searched against the TAIR database. Trypsin with a maximum of 3 missed cleavages and one unspecific end was selected as the protease. Carbamidomethylation (Cys) was set as a fixed modification, and oxidation (Methionine) and deamination (Asparagine, Glutamine) were set as variable modifications. Precursor mass tolerance was set as 15 ppm. Fragment mass tolerances for CID were set to 0.8 Da, respectively. All spectra were manually validated. For all peptides present at −10*Log(P) > 20, considered as high-confidence, spectra were manually checked, validated, or disqualified.

## Acknowledgements

The BioDiaAlk probe was kindly provided by Prof. Kate Carroll (Florida Atlantic University, USA).

## Author contributions

Y.T., S.A., P.P., E.F. and B.G. conceptualized and designed the study. Y.T., S.A., A.A., L.P., Z.D., G.N., M.L-P., D.M.G., N.L.M. and S.L. performed the experiments. Y.T., S.A., A.A., M.L-P., E.F. and B.G contributed to data analysis and figure preparation. P.P., C.G., M.S., E.F. and B.G. provided funding acquisition for the study. Y.T., S.A., E.F., and B.G. wrote the manuscript with critical input from all authors.

