## Supplemental Figures and Tables for "Hydrogen sulfide modulates plant hypoxic responses through the persulfidation of Plant Cysteine Oxidases"

**This PDF file includes:**

Suppl. Figures 1-12

Suppl. Tables 1-2

SI References

**Supplementary figures**

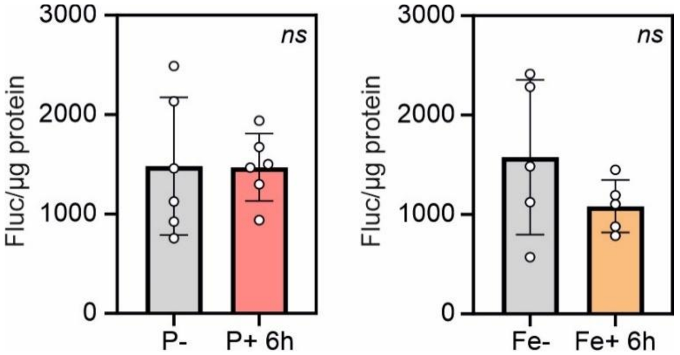


**Suppl. Figure 1. Impact of nutrient re-supply on RAP2.12$_{1-28}$ stability in *28RAPFluc*. b)** RAP2.12$_{1-28}$ stability after 5 days of growth in phosphorous (P- = 25 µM P), or **c)** iron (Fe- = 0 µM Fe) liquid deficiency medium and after 6 hours of recovery in complete nutrient liquid media (mean ± SD; n=5). Relative luciferase activity was normalised on total proteins. Student's t-test was used to investigate the statistical differences between treatments (*ns*, not significant). For detailed nutrient media composition see **Suppl. Table 1.**

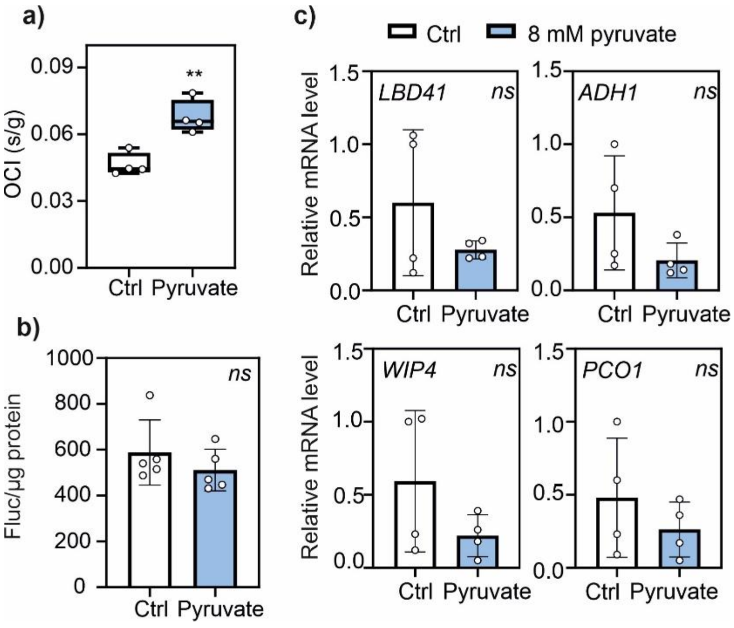


**Suppl. Figure 2. Effect of pyruvate treatment on $O_2$ consumption and anaerobic responses. a)** $O_2$ consumption in 5 day-old Col-0 seedlings treated with 8 mM pyruvate for 6 hours. OCI, Oxygen Consumption Index. Data are mean ± SD (n=4). **b)** Anaerobic marker gene expression in the same samples as (a) (mean ± SD; n=4). **c)** *28RAPFluc* seedlings were grown for five days in liquid media and treated with 8 mM pyruvate. Luciferase activity was analysed after 6 hours from the pyruvate supplementation and normalised on total proteins. Asterisk indicate significantly differences between treatments after Student's t-test (*p<0.05; ** p<0.01, *** p<0.001), *ns* stands for not significant.

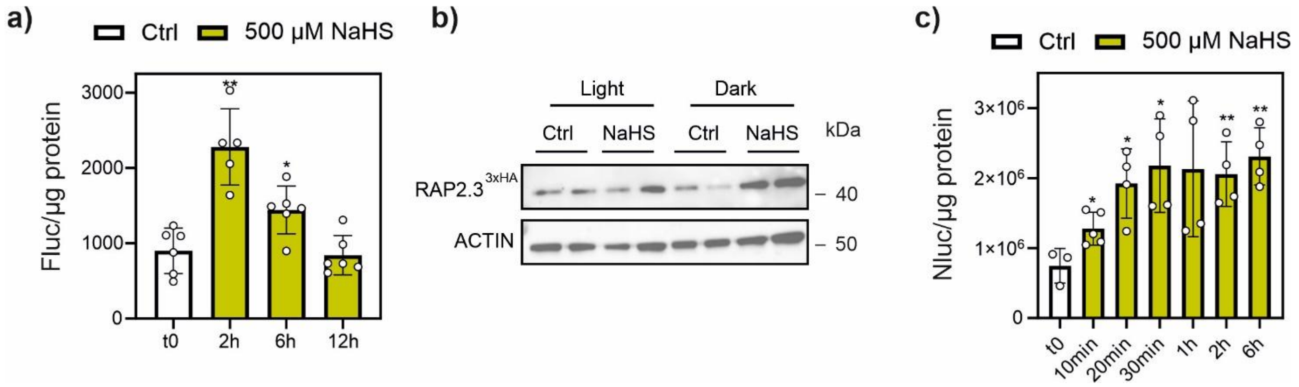


**Suppl. Figure 3. Impact of elevated $H_2S$ supplementation on anaerobic responses. a)** Time course detection of firefly luciferase activity in 7 day-old *28RAPFluc* seedlings exposed to 500 µM NaHS. Data are represented as mean ± SD (n=6). Activity data were normalized on extract total protein content. **b)** Immunoblotting of RAP2.3-HA protein in *35S:RAP2.3*$^{3xHA}$ seedlings treated with 500 µM NaHS for 2 hours in dark and light conditions. Equal loading was assessed by blotting of the Actin-11 housekeeping protein. Two biological replicates are shown. **c)** Time course detection of Nanoluciferase activity in *HRPE:Nluc* seedlings after the treatment with 500 µM NaHS. Activity data were normalized on extract total protein content. Asterisks in (a) and (c) indicate statistically significant differences between treated and untreated ($t_0$) samples after Student's t-test (* p<0.05; ** p<0.01; *** p<0.001).

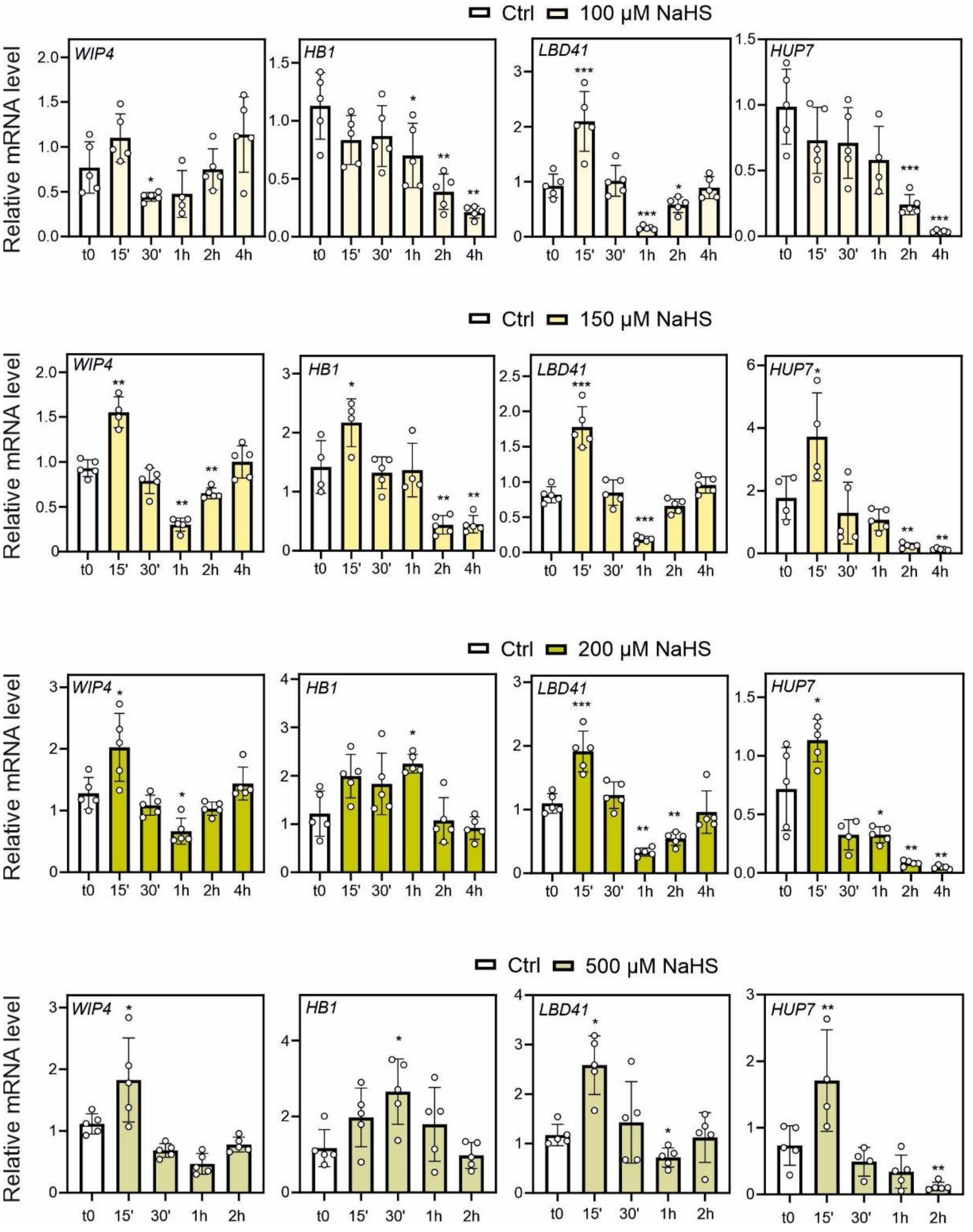


**Suppl. Figure 4. Anaerobic genes rapidly induced after different $H_2S$ supplementation.** Effect of 100, 150, 200 and 500 μM $H_2S$ supplementation on *WIP4*, *HB1*, *LBD41* and *HUP7* expression. Asterisks indicate statistically significant differences between treated and t0 samples after Student's t-test (* $p<0.05$; ** $p<0.01$; *** $p<0.001$).

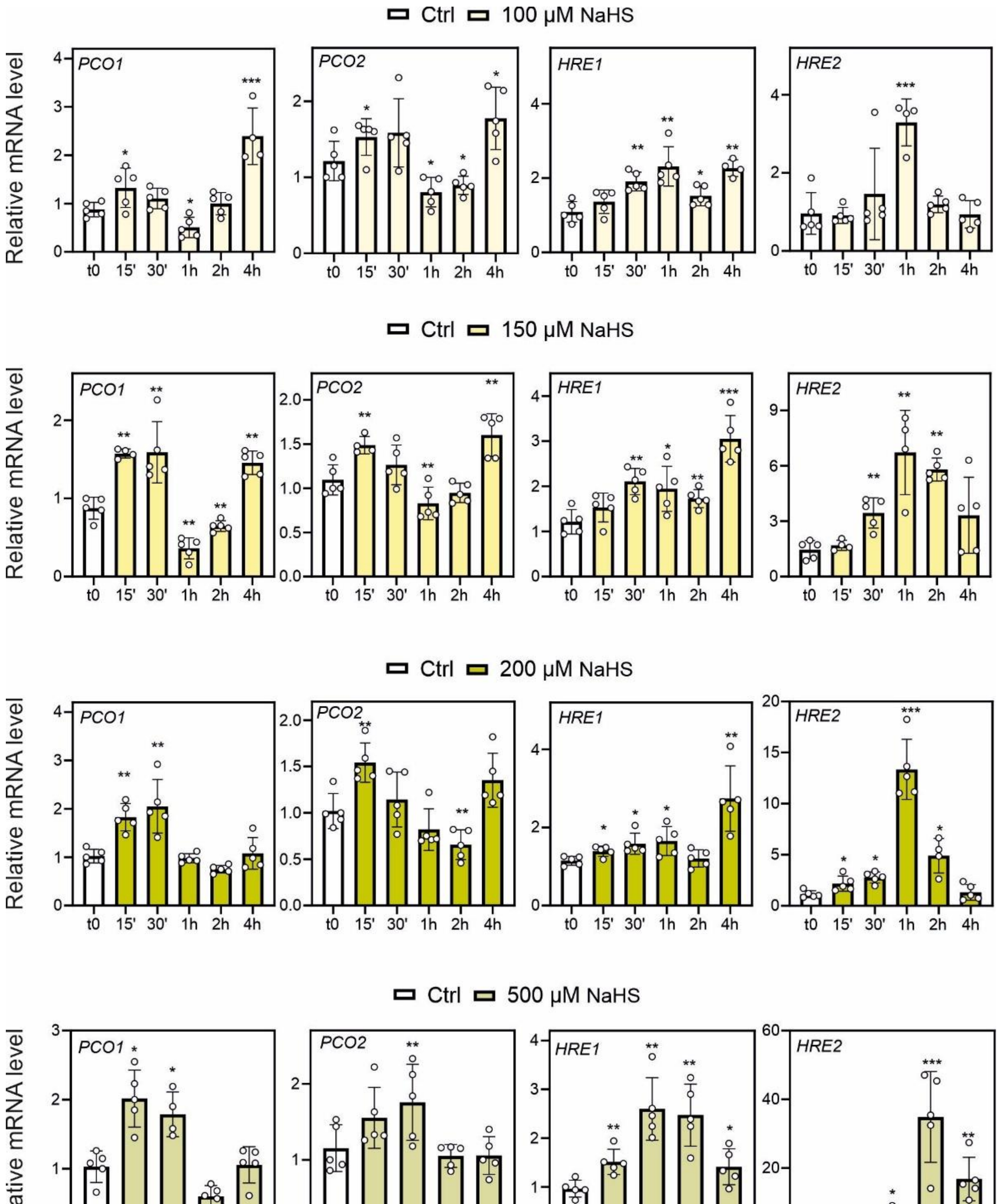


**Suppl. Figure 5. Anaerobic genes induced stably or slowly after different $H_2S$ supplementation.** Effect of $H_2S$ supplementation at 100, 150, 200, and 500 μM on the expression of *PCO1*, *PCO2*, *HRE1*, and *HRE2*. Asterisks indicate statistically significant differences compared to t0, as determined by Student's t-test (*$p < 0.05$; **$p < 0.01$; **$p < 0.001$).

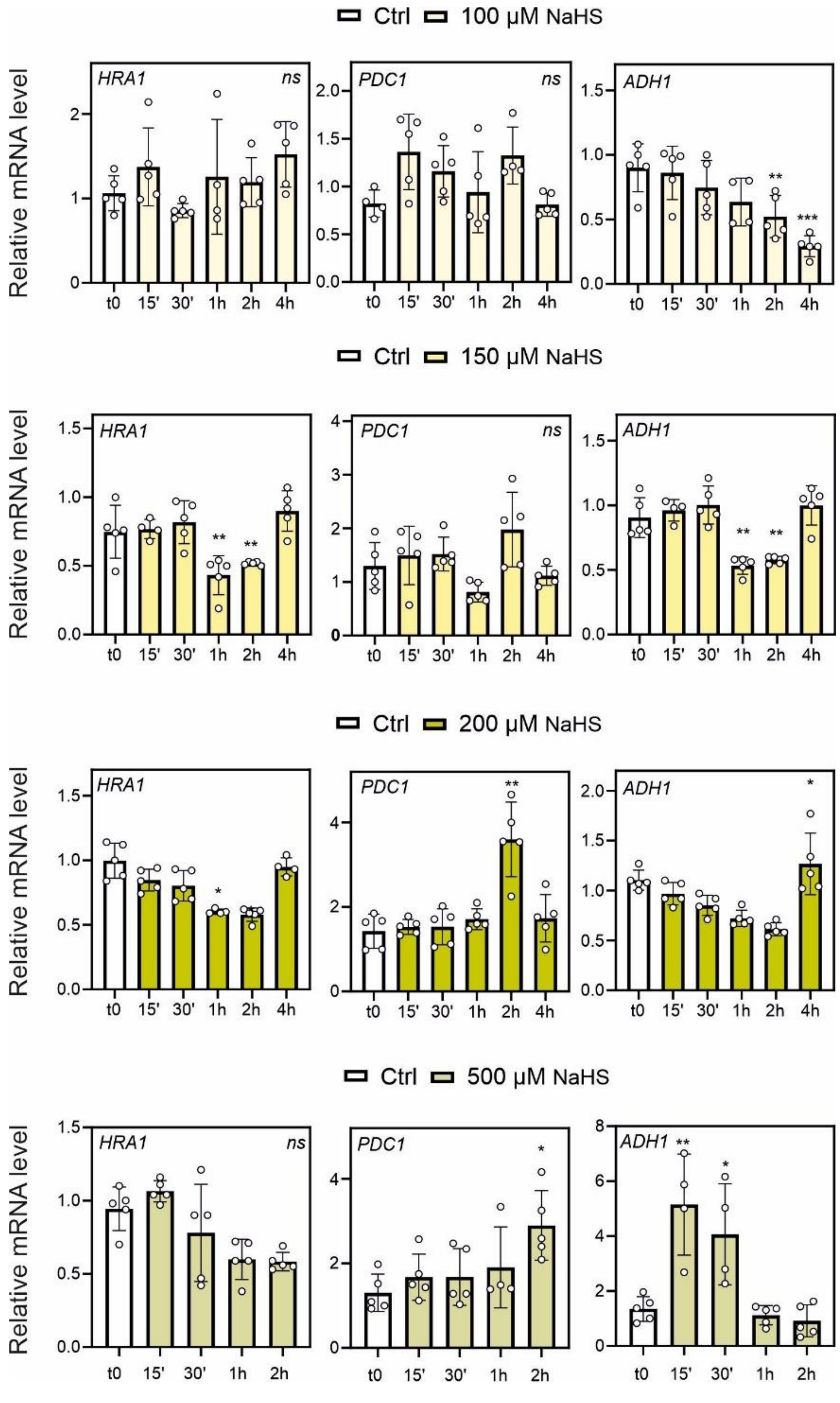


**Suppl. Figure 6. Anaerobic genes weakly induced after different $H_2S$ supplementation.** *HRA1*, *PDC1*, and *ADH1* expression in response to $H_2S$ supplementation at 100, 150, 200, and 500 µM. Statistically significant differences from t0 are indicated by asterisks, based on Student's t-test (*p < 0.05; **p < 0.01; **p < 0.001).

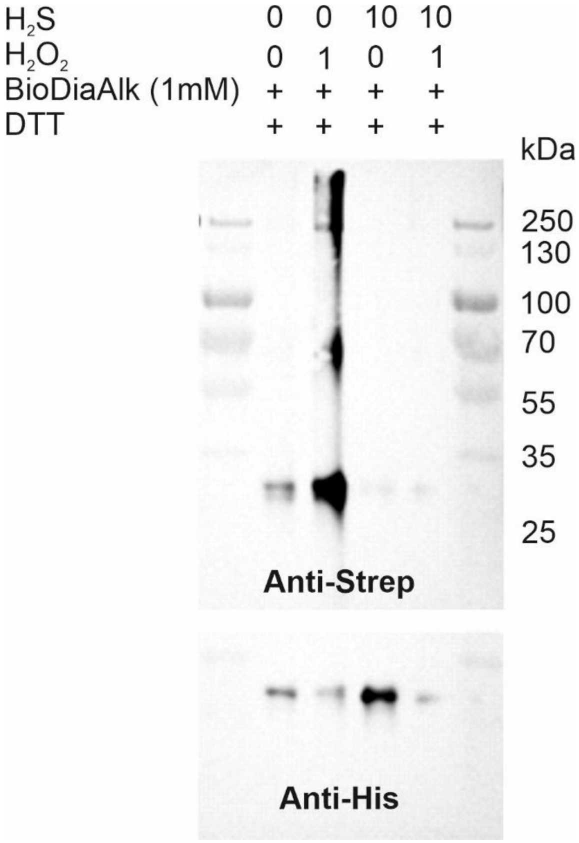


**Suppl. Figure 7. Supporting blot image.** Full-size blot image for Fig. 4a.

a)

```
CLUSTAL multiple sequence alignment by MUSCLE (3.8)
PCO3            MLSRLFKAGEKVLSNLVSKKDIYMASRNQEKSPK-------------------------V
PCO4            M--PYF-----------------------------------------------------A
PCO5            M--PYF-----------------------------------------------------I
PCO1            M—GFEMKPEKEVLELISSKNQCKSNPNSVKKKNKNKN-KKMMMTWRRKKIDSPADGITAV
PCO2            M------GTDTVMSGRVRKDLSKTNPNGNIPENRSNSRKKIQ--RRSKKTLICP-----V

PCO3            QELYDLCKETFTGKAPS--PASMAIQKLCSVLDSVSPADVGLEEVSQDDDRGYGVSGVSR
PCO4            QRLYNTCKASFSSDGP---ITEDALEKVRNVLEKIKPSDVGIEQDAQLARSRSGP--LNE
PCO5            QRLFNTCKSSLSPNGP---VSEEALDKVRNVLEKIKPSDVGLEQEAQLVRNWPGP--GNE
PCO1            RRLFNTCKEVFSNGGPGVIPSEDKIQQLREILDDMKPEDVGLTPTMPYFRPNSGV-----
PCO2            QKLFDTCKKVFADGKSGTVPSQENIEMLRAVLDEIKPEDVGVNPKMSYFRST--------
                . *:: **  ::   .    :.  :: :  :*:.:.* ***:
PCO3            FNRVGRWAQPITFLDIHECDTFTMCIFCFPTSSVIPLHDHPEMAVFSKILYGSLHVKAYD
PCO4            RNGSNQSPPAIKYLHLHECDSFSIGIFCMPPSSMIPLHNHPGMTVLSKLVYGSMHVKSYD
PCO5            RNGNHHSLPAIKYLQLHECDSFSIGIFCMPPGSIIPLHNHPGMTVLSKLVYGSMHVKSYD
PCO1            ---EARSSPPITYLHLHQCDQFSIGIFCLPPSGVIPLHNHPGMTVFSKLLFGTMHIKSYD
PCO2            ---VTGRSPLVTYLHIYACHRFSICIFCLPPSGVIPLHNHPEMTVFSKLLFGTMHIKSYD
                          :.:* :: *  *:: ***:*...:****:** *:*:**:::*::*:*:**
PCO3            WVEPPCIITQDKGVPGSLPARLAKLVSDKVITPQSEIPALYPKTGGNLHCFTALTPCAVL
PCO4            WLEPQLTEPEDP----SQEARPAKLVKDTEMTAQSPVTTLYPKSGGNIHCFKAITHCAIL
PCO5            WAEPDQSELDDP-----LQARPAKLVKDIDMTSPSPATTLYPTTGGNIHCFKAITHCAIF
PCO1            WV-VDAPMRDSK-------TRLAKLKVDSTFTAPCNASILYPEDGGNMHRFTAITACAVL
PCO2            WV-PDSPQPSSD-------TRLAKVKVDSDFTAPCDTSILYPADGGNMHCFTAKTACAVL
                *        ..        :* **:  *  :*. .  . ***  ***:* *.* * **::
PCO3            DILSPPYKESVGRSCSYYMDYPFSTFALENGMKKV-DEGKEDEYAWLVQI-DTPDDLHMR
PCO4            DILAPPYSSEHDRHCTYFRKSRREDLP---GELEV-DGEVVTDVTWLEEF-QPPDDFVIR
PCO5            DILSPPYSSTHGRHCNYFRKSPMLDLP---GEIEVMNGEVISNVTWLEEY-QPPDNFVIW
PCO1            DVLGPPYCNPEGRHCTYFLEFPLDKLSSEDDDVLS-SEEEKEGYAWLQERDDNPEDHTNV
PCO2            DVIGPPYSDPAGRHCTYYFDYPFSSFSV--DGVVV-AEEEKEGYAWLKEREEKPEDLTVT
                *::.*** .  .* *.*: .     :.   .             :** :  : *::
PCO3            PGS-YTGPTIRV
PCO4            RIP-YRGPVIRT   Cys residues conserved in all PCOs
PCO5            RVP-YRGPVIRK   Cys residues conserved in all PCOs except PCO1
PCO1            VGALYRGPKVED
PCO2            ALM-YSGPTIKE
                    * ** :
```

b)

| # | b | Seq | y | # |
|---|---|---|---|---|
| 1 | 72.04 | A | | 21 |
| 2 | 185.13 | I | 2340.12 | 20 |
| 3 | 286.18 | T | 2227.03 | 19 |
| 4 | 423.24 | H | 2125.99 | 18 |
| 5 | 615.24 | C(+89.00) | 1988.93 | 17 |
| 6 | 686.28 | A | 1796.92 | 16 |
| 7 | 799.36 | I | 1725.88 | 15 |
| 8 | 912.44 | L | 1612.80 | 14 |
| 9 | 1027.47 | D | 1499.71 | 13 |
| 10 | 1140.55 | I | 1384.69 | 12 |
| 11 | 1253.65 | L | 1271.60 | 11 |
| 12 | 1324.68 | A | 1158.52 | 10 |
| 13 | 1421.73 | P | 1087.48 | 9 |
| 14 | 1518.79 | P | 990.43 | 8 |
| 15 | 1681.85 | Y | 893.38 | 7 |
| 16 | 1768.88 | S | 730.31 | 6 |
| 17 | 1855.92 | S | 643.28 | 5 |
| 18 | 1984.96 | E | 556.25 | 4 |
| 29 | 2122.02 | H | 427.21 | 3 |
| 20 | 2237.04 | D | 290.15 | 2 |
| 21 | | R | 175.12 | 1 |

c)

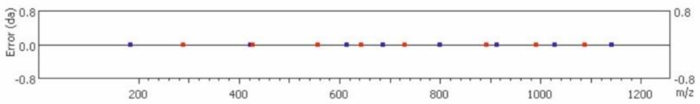


**Suppl. Figure 8. Investigation of PCO4 PTMs. a)** Multiple sequence alignment of PCO proteins performed using MUSCLE via ClustalOmega. Cys residues highlighted in yellow are conserved in all PCOS members (referred to PCO4 sequence: Cys$^{12}$, Cys$^{79}$, Cys$^{88}$, Cys$^{172}$, and Cys$^{190}$). PCO4 Cys$^{165}$ residue highlighted in blue is conserved in all PCOs with the exception of PCO1. **b)** Table of theoretical and observed fragment ion masses for a representative PCO4 peptide containing the S–S–IAM modification on Cys172. Expected b- and y-ion series are listed, with matched ions highlighted in blue (b-series) and red (y-series). **c)** The "Error Map" shows the mass accuracy assessment of identified fragment ions. The x-axis represents the m/z values of matched fragments, and the y-axis shows the deviation in Daltons. Each dot corresponds to a matched ion, indicating high mass accuracy for the assigned peaks.

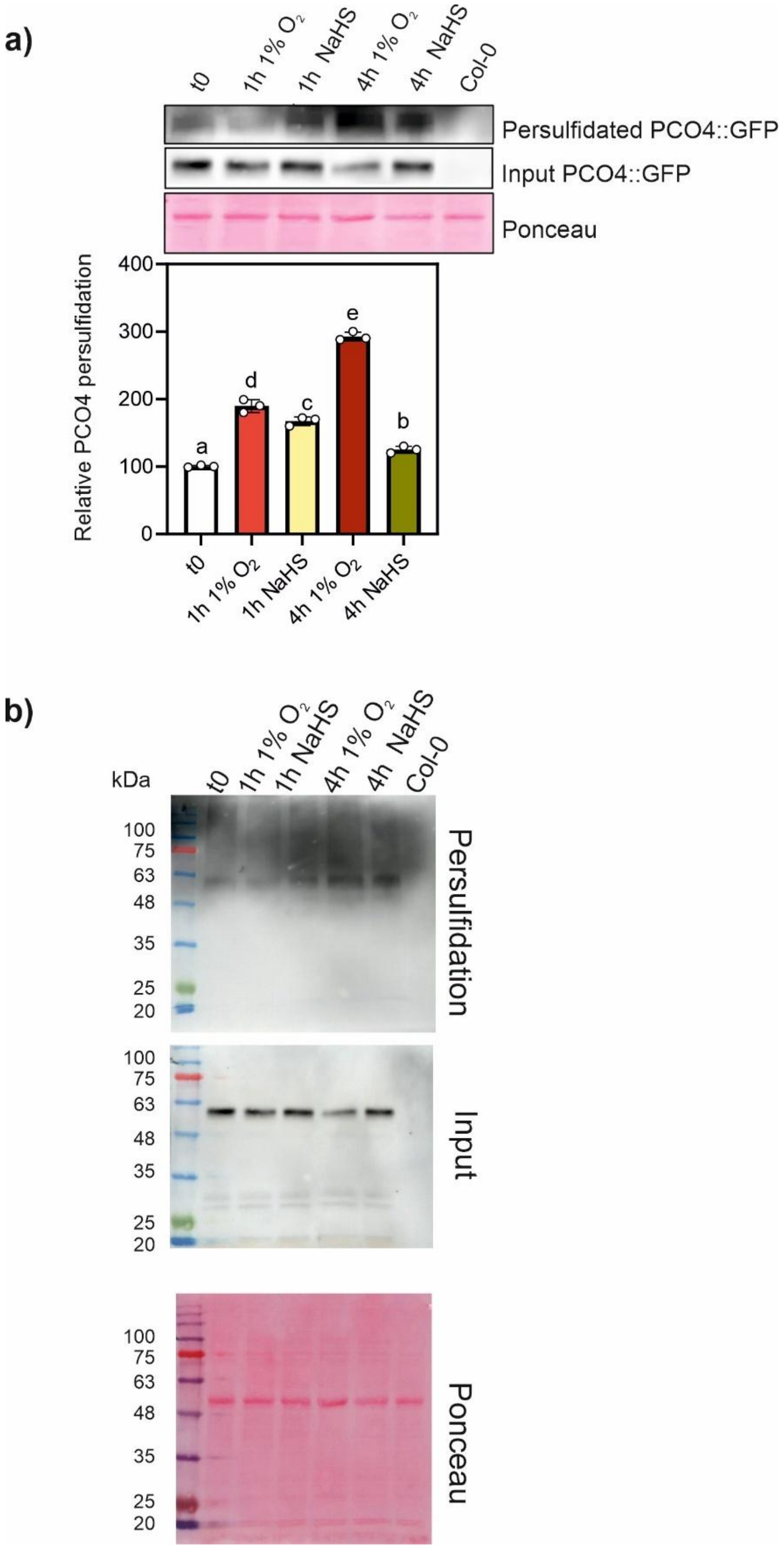


**Suppl. Figure 9. Supporting blot images. a)** Immunoblot and *in vivo* relative quantification of PCO4 persulfidation after 1 h or 4 h of low oxygen treatment or 500 µM $H_2S$ exposure in *pPCO4:PCO4:GFP* seedlings in the *4pco* background. PCO4 persulfidation levels are shown relative to PCO4 input. Col-0 was used as negative control. Letters indicate significant differences (one-way ANOVA, Tukey post-hoc). **b)** Uncropped images for Suppl. Figure. 9a.

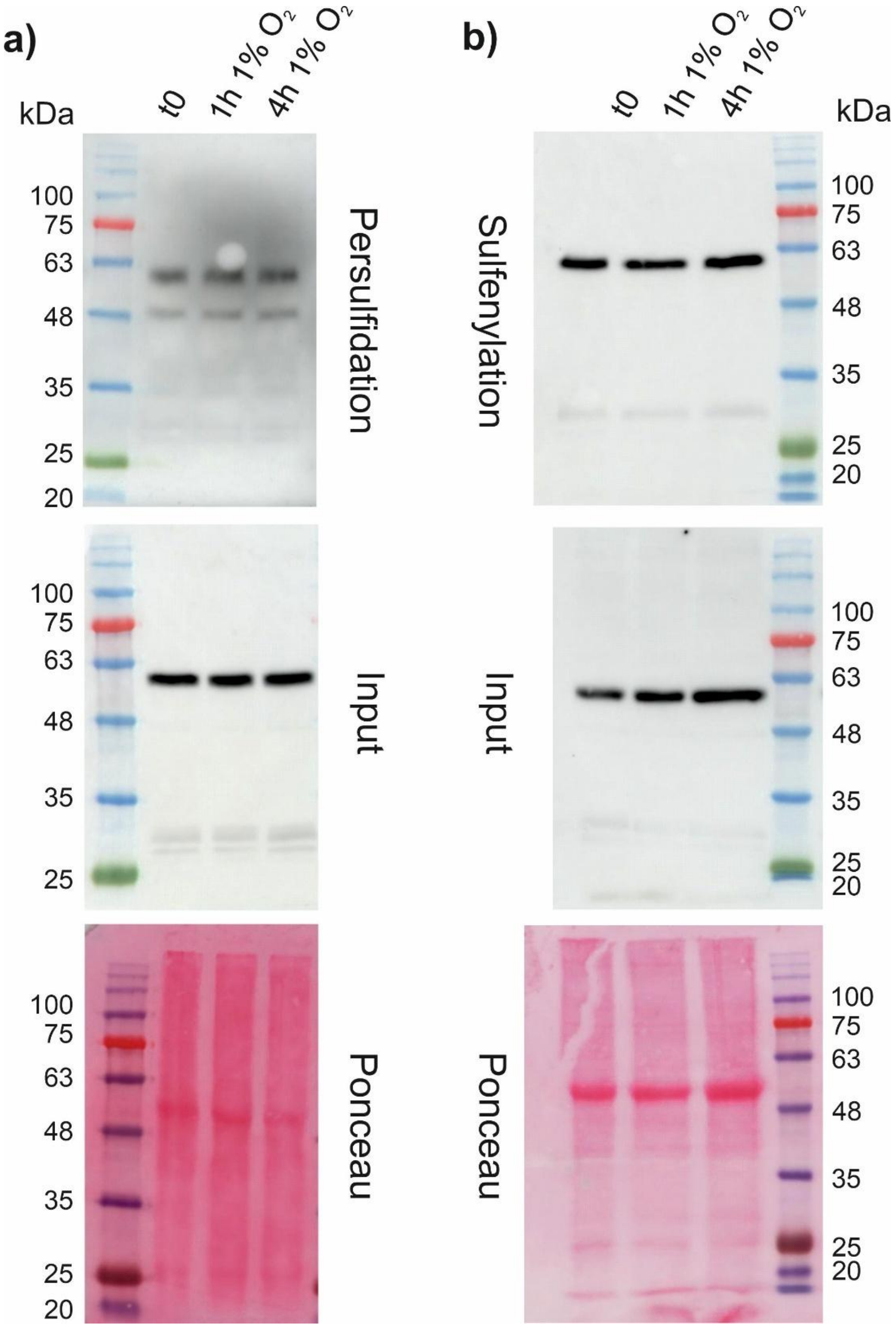


**Suppl. Figure 10. Supporting blot images. a)** Uncropped images for Fig. 5a and **b)** Fig. 5d.

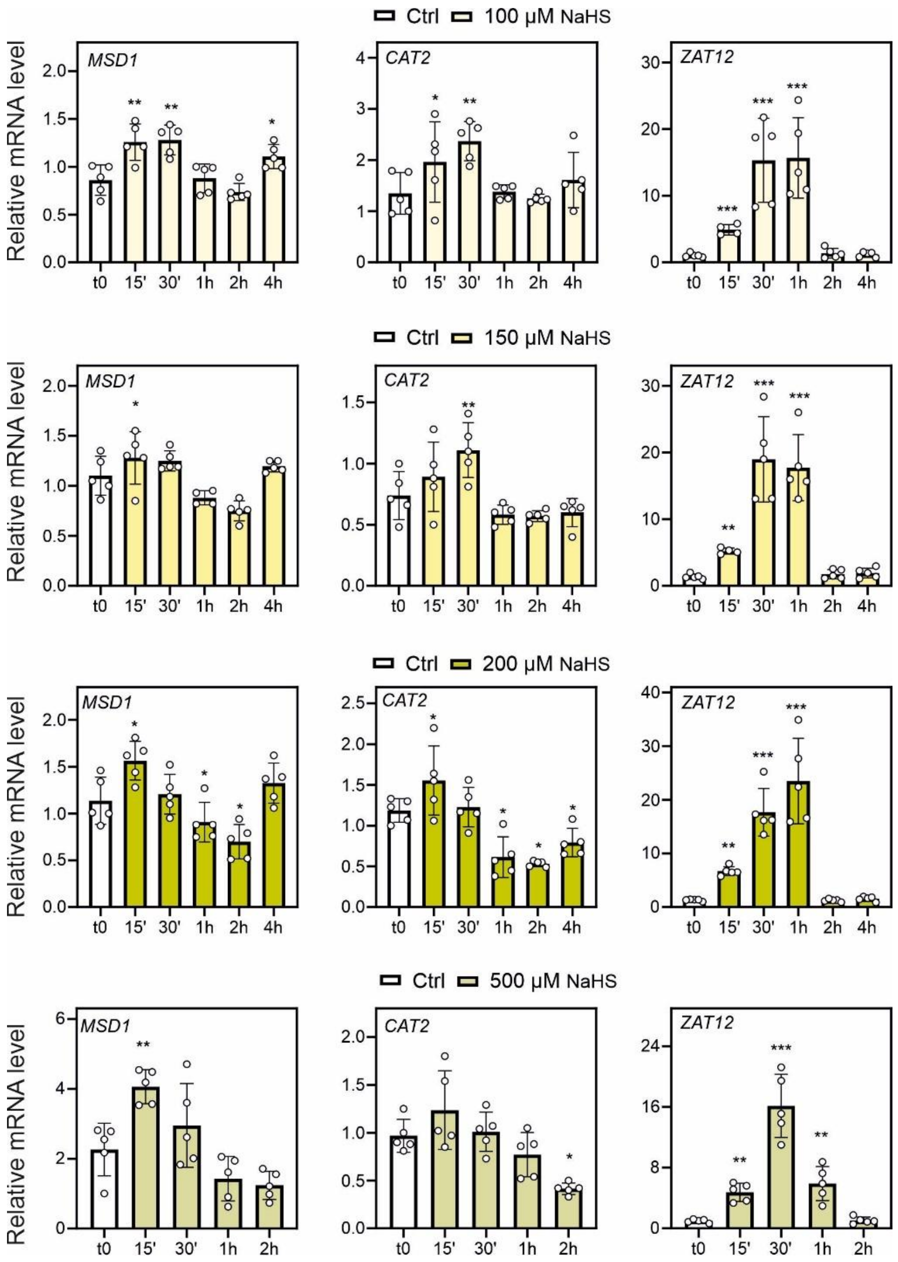


**Suppl. Figure 11. Effect of different $H_2S$ supplementation on ROS responsive genes expression.** Impact of $H_2S$ supplementation at concentrations of 100, 150, 200, and 500 µM on the expression levels of *MSD1, CAT2* and *ZAT12*. Asterisks denote statistically significant differences between treated samples and t0, as determined by Student's t-test (* $p<0.05$; ** $p<0.01$; *** $p<0.001$).

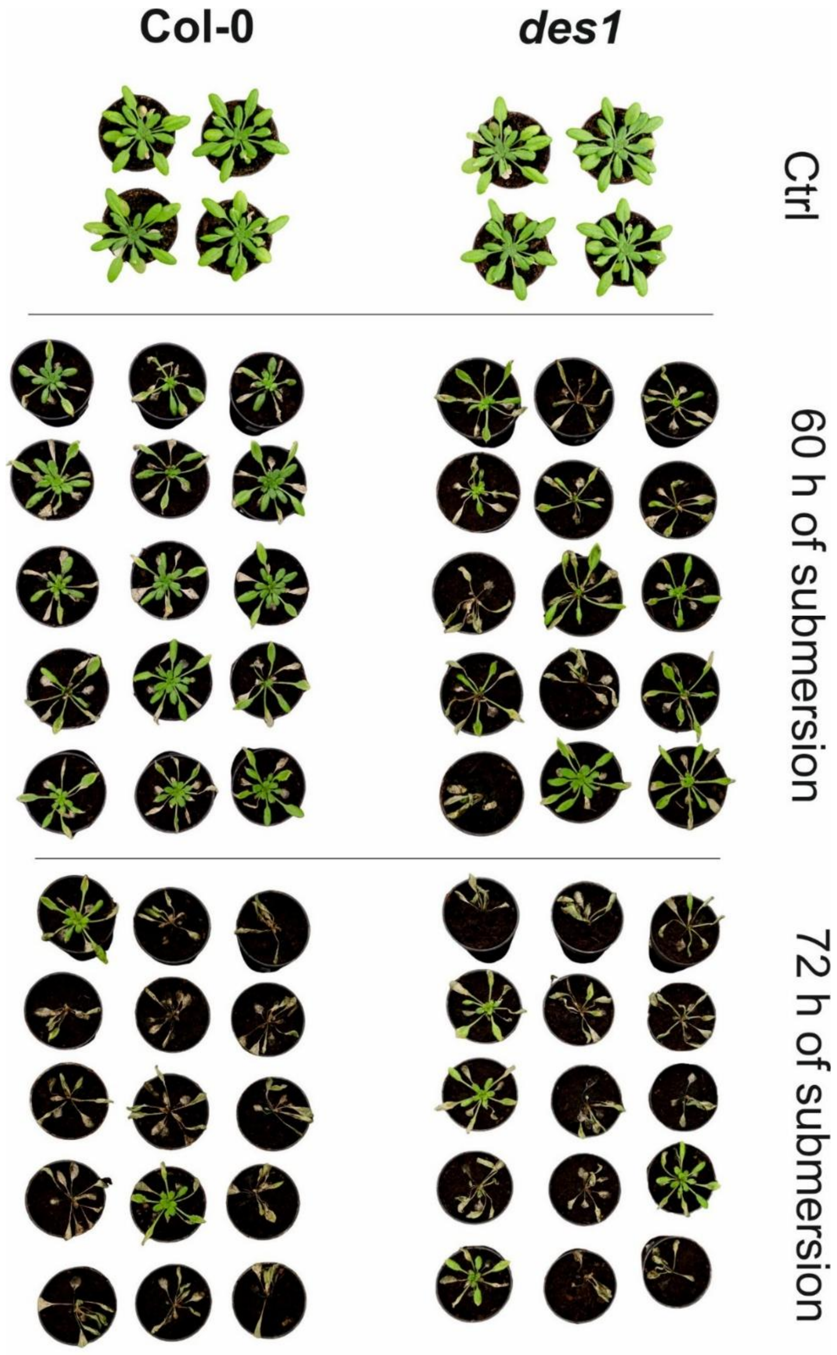


**Suppl. Figure 12. Images of Col-0 and *des1* plants exposed to submergence.** 21-day-old Col-0 and *des1* plants one week after dark submergence for 60 or 72 hours. Control plants were kept in the dark for 60 hours without submergence.

### Supplemental tables

**Suppl. Table 1. Meta-analysis of *HRG* expression from the microarray study in Bielcka et al. (2015).** Expression of 49 core HRGs (Mustroph *et al.,* 2010) in seedlings after 30 minutes (30') or 3 hours from S re-integration. Data are expressed as fold change (FC) relative to S deficiency (S-, 150 µM).

| | | **Experiment 1** | | **Experiment 2** | |
|---|---|---|---|---|---|
| Gene name | AGI Code | FC 30'/S- | FC 3h/S- | FC 30'/S- | FC 3h/S- |
| *HRA1* | *AT3G10040* | 2,91 | 1,47 | 1,44 | 1,49 |
| *AT5G26200* | *AT5G26200* | 2,33 | 1,67 | 0,25 | 0,50 |
| *ATPP2-A11* | *AT1G63090* | 2,06 | 1,11 | 1,50 | 1,15 |
| *PCO1* | *AT5G15120* | 1,72 | 0,94 | 1,18 | 0,96 |
| *PDC1* | *AT4G33070* | 1,69 | 1,56 | 3,65 | 2,23 |
| *CML38* | *AT1G76650* | 1,67 | 0,21 | 0,97 | 1,38 |
| *WIP4* | *AT4G10270* | 1,65 | 1,15 | 0,86 | 1,29 |
| *CYP707A3* | *AT5G45340* | 1,65 | 1,02 | 1,44 | 0,96 |
| *HUP36* | *AT1G19530* | 1,58 | 2,15 | 0,86 | 1,04 |
| *HUP39* | *AT3G23170* | 1,55 | 0,87 | 1,31 | 0,74 |
| *AT5G02200* | *AT5G02200* | 1,48 | 0,81 | 1,07 | 0,87 |
| *SRO5* | *AT5G62520* | 1,42 | 0,34 | 2,29 | 0,98 |
| *NIP2.1* | *AT2G34390* | 1,42 | 1,69 | 0,38 | 0,60 |
| *HUP40* | *AT4G24110* | 1,41 | 0,91 | 1,05 | 1,11 |
| *AT1G72940* | *AT1G72940* | 1,38 | 1,02 | 1,29 | 1,40 |
| *WIP5* | *AT4G33560* | 1,37 | 1,76 | 1,59 | 1,37 |
| *PHI-1* | *AT1G35140* | 1,33 | 1,09 | 0,86 | 0,65 |
| *HUP9* | *AT5G10040* | 1,29 | 1,57 | 1,03 | 2,03 |
| *ATL23* | *AT5G42200* | 1,26 | 0,72 | 0,91 | 1,00 |
| *PDC2* | *AT5G54960* | 1,23 | 0,78 | 1,08 | 0,83 |
| *ACR7* | *AT4G22780* | 1,19 | 0,63 | 0,95 | 0,62 |
| *HUP54* | *AT4G27450* | 1,15 | 1,00 | 0,98 | 1,53 |
| *JAI3* | *AT3G17860* | 1,15 | 0,99 | 1,08 | 0,95 |
| *HRE2* | *AT2G47520* | 1,12 | 1,46 | 0,88 | 5,22 |
| *AT5G47060* | *AT5G47060* | 1,07 | 0,79 | 1,13 | 0,74 |
| *ACHT5* | *AT5G61440* | 1,04 | 1,01 | 0,88 | 1,32 |
| *ATPP2-A13* | *AT3G61060* | 1,01 | 1,50 | 0,81 | 1,22 |
| *HUP42* | *AT4G39675* | 1,00 | 5,00 | 0,50 | 1,49 |
| *RBOHD* | *AT5G47910* | 0,99 | 0,68 | 1,24 | 0,80 |
| *FLZ13* | *AT1G74940* | 0,99 | 1,10 | 1,06 | 0,98 |
| *ADH1* | *AT1G77120* | 0,98 | 1,31 | 1,18 | 1,98 |
| *PCO2* | *AT5G39890* | 0,96 | 1,33 | 1,40 | 1,47 |
| *AT1G26270* | *AT1G26270* | 0,95 | 0,87 | 1,00 | 0,79 |
| *AT5G44730* | *AT5G44730* | 0,95 | 1,05 | 1,16 | 1,19 |
| *ALAAT1* | *AT1G17290* | 0,94 | 1,22 | 1,04 | 1,28 |
| *TIL* | *AT5G58070* | 0,92 | 0,96 | 0,87 | 0,74 |
| *HUP6* | *AT3G27220* | 0,91 | 1,10 | 1,11 | 1,20 |
| *SUS4* | *AT3G43190* | 0,90 | 0,82 | 1,28 | 1,11 |

| | | | | | |
|---|---|---|---|---|---|
| *AT1G55810* | *AT1G55810* | 0,89 | 1,31 | 1,01 | 0,96 |
| *PFK6* | *AT4G32840* | 0,89 | 1,18 | 0,94 | 1,07 |
| *ETR2* | *AT3G23150* | 0,87 | 0,60 | 1,07 | 0,76 |
| *AT2G17850* | *AT2G17850* | 0,86 | 0,71 | 1,35 | 1,63 |
| *AT4G17670* | *AT4G17670* | 0,79 | 0,42 | 0,97 | 0,51 |
| *PGB1* | *AT2G16060* | 0,79 | 1,68 | 1,55 | 2,74 |
| *LBD41* | *AT3G02550* | 0,76 | 0,72 | 0,90 | 0,90 |
| *ACO1* | *AT2G19590* | 0,73 | 0,74 | 1,36 | 1,20 |
| *FTM1* | *AT1G43800* | 0,62 | 0,20 | 1,28 | 1,17 |
| *HUP44* | *AT5G66985* | 0,49 | 1,63 | 1,37 | 1,68 |
| *HUP32* | *AT1G33055* | 0,25 | 0,84 | 0,93 | 1,76 |

**Suppl. Table 2. Nutrient media composition.**

| Nutrient | 25 µM sulfur (S-) | 0 µM iron (Fe-) | 25 µM phosphorus (P-) | Ctrl |
|---|---|---|---|---|
| | Final concentration µM | Final concentration µM | Final concentration µM | Final concentration µM |
| $KH_2PO_4$ | 625 | 625 | 25 | 625 |
| $NH_4NO_3$ | 1000 | 1000 | 1000 | 1000 |
| $KNO_3$ | 9400 | 9400 | 10000 | 9400 |
| $CaCl_2$ $2H_2O$ | 1500 | 1500 | 1500 | 1500 |
| MES pH 5.5 with KOH | 1000 | 1000 | 1000 | 1000 |
| $H_3BO_3$ | 50 | 50 | 50 | 50 |
| KI | 2.5 | 2.5 | 2.5 | 2.5 |
| $MnCl_2$ $4H_2O$ | 50 | 50 | 50 | 50 |
| $ZnCl_2$ | 15 | 0 | 0 | 15 |
| NaFe EDTA | 75 | 0 | 75 | 75 |
| $CoCl_2$ $6H_2O$ | 0.055 | 0.055 | 0.055 | 0.055 |
| $CuCl_2$ $4H_2O$ | 0.053 | 0.053 | 0.053 | 0.053 |
| $Na_2MoO_4$ $2H_2O$ | 0.52 | 0.52 | 0.52 | 0.52 |
| $MgSO_4$ $7H_2O$ | 25 | 25 | 25 | 750 |
| $MgCl_2$ $6H_2O$ | 725 | 0 | 0 | 0 |

**Suppl. Table 3. RT-PCR primers used in this study.**

| Gene | AGI code | Forward primer sequence | Reverse primer sequence |
|---|---|---|---|
| *SULTR1;1* | *AT4G08620* | AGATCGGTCTCTTGATCGCTGTG | AACCGTGGTTCTTGGTCTCGTC |
| *SULTR4;2* | *AT3G12520* | CGCTCACAGGTCTTGCTCTT | AGCCGATGTTGGAAGCAGTA |
| *ADH1* | *AT1G77120* | TATTCGATGCAAAGCTGCTGTG | CGAACTTCGTGTTTCTGCGGT |
| *PCO1* | *AT5G15120* | ATTGGGTGGTTGATGCTCCAATG | ATGCATGTTCCCGCCATCTTC |
| *LBD41* | *AT3G02550* | TGAAGCGCAAGCTAACGCA | ATCCCAGGACGAAGGTGATTG |
| *PDC1* | *AT4G33070* | CACAGAATCTTCAATGCTTCTTACC | CCATGATAAAGCGTACATGGAA |
| *WIP4* | *AT4G10270* | ATGGACAGTGGCAGTGAGCATC | GTTCCACCGACAAAGACCTAGTTG |
| *HUP7* | *AT1G43800* | TTGGCAACCCGCTTCTTTCTTACC | TTTCCCTCAGCTCACGAACCTG |
| *HRE2* | *AT2G47520* | GAAGCGTAAACCCGTCTCAGTG | TTTGCTCGGGTCACGAATCT |
| *HRA1* | *AT3G10040* | TCATGTTACGGCGGAGTGAA | CAACCCGTGTACCCGAAGAC |
| *HB1* | *AT2G16060* | TTTGAGGTGGCCAAGTATGCA | TGATCATAAGCCTGACCCCAA |
